# Axial-circular magnetic levitation assisted biofabrication and manipulation of cellular structures

**DOI:** 10.1101/2021.01.26.428192

**Authors:** Muge Anil-Inevi, Kerem Delikoyun, Gulistan Mese, H. Cumhur Tekin, Engin Ozcivici

## Abstract

Diamagnetic levitation is an emerging technology for remote manipulation of cells in cell and tissue level applications. Low-cost magnetic levitation configurations using permanent magnets are commonly composed of a culture chamber physically sandwiched between two block magnets that limit working volume and applicability. This work describes a single ring magnet-based magnetic levitation system to eliminate physical limitations for biofabrication. Developed configuration utilizes sample culture volume for construct size manipulation and long-term maintenance. Furthermore, our configuration enables convenient transfer of liquid or solid phases during the levitation. Prior to biofabrication, we first calibrated the platform for levitation with polymeric beads, considering the single cell density range of viable cells. By taking advantage of magnetic focusing and cellular self-assembly, millimeter-sized 3D structures were formed and maintained in the system allowing easy and on-site intervention in cell culture with an open operational space. We demonstrated that the levitation protocol could be adapted for levitation of various cell types (i.e., stem cell, adipocyte and cancer cell) representing cells of different densities by modifying the paramagnetic ion concentration that could be also reduced by manipulating the density of the medium. This technique allowed the manipulation and merging of separately formed 3D biological units, as well as the hybrid biofabrication with biopolymers. In conclusion, we believe that this platform will serve as an important tool in broad fields such as bottom-up tissue engineering, drug discovery and developmental biology.

## Introduction

Magnetic force-based manipulation of the living cells has emerged as a powerful tool for cellular and tissue level bioengineering applications [1–4]. With the advances in technology and the improvements in design, magnetic manipulation systems with different complexity have been developed for various biotechnological goals including isolation and enrichment of rare cells [5, 6] and guiding cells into a particular spatial arrangement in 2 dimensional (2D) or 3 dimensional (3D) cultures [7–9]. Compared to the alternative operation principles such as electrical, optical and acoustic force-based techniques, magnetic manipulation offers various advantages such as minimal impact on cell viability, simple and low-cost design, and low sensitivity to environmental parameters such as ionic concentration and pH [10].

Cell magnetophoresis can be performed in two ways, either by manipulating the magnetic susceptibility of the cells or manipulating the magnetic susceptibility of the environment that cells are found [1–3]. Cells, that exhibit greater magnetic susceptibility than their surrounding buffer or medium due to labeling with magnetic particles or a rare intrinsic property of some cell types (i.e. paramagnetic hemoglobin containing blood cells and magnetotactic bacteria), move towards regions of the high magnetic field (positive magnetophoresis) [11]. However, most cell types are diamagnetic in nature, and once placed into a surrounding environment with high magnetic susceptibility, they are repelled towards the minimal magnetic field (negative magnetophoresis, also referred to as diamagnetophoresis) [12–17]. Stable cell trapping and self-assembly have been previously conducted by both positive and negative magnetophoresis to create viable 3D structures [18–20]. In positive magnetophoresis, it is possible to manipulate cells even with extremely small magnetic gradients using magnetic labels, allowing cell culture to reach high volume ratios up to several milliliters [21, 22]. However, this manipulation technique is challenging because labelling process is time-consuming and manually intensive [23] as well as prone to experimental variability based on variations in magnetic moments of beads [24] or cell labeling efficiency [25].

Negative magnetophoresis-based magnetic levitation of cells in contrast, benefits from a label-free methodology. This approach was conventionally performed under a strong magnetic field generated by electromagnets due to the low magnetic susceptibility difference between the biological material and its surroundings [26]. As a simple and low-cost alternative, permanent magnets have been recently used to levitate diamagnetic objects in paramagnetic salt solutions or ferrofluids under weak magnetic fields [3, 27, 28]. A magnetic levitation configuration was proposed to levitate diamagnetic objects based on their physical properties. This system levitates materials in paramagnetic solutions under a low magnetic field (<0.5 T) that is generated by two rectangular permanent magnets with the same poles facing each other [13, 15, 29]. However, this setup only allows biofabrication applications in microcapillaries, limiting working volumes for cells [19, 20, 30, 31]. Increasing the size of living structures is of prime importance for straightforward implementation of testing protocols by providing an adequate number of cells [32–34], and for the production of sizable tissue engineering constructs [35, 36]. A technique has been previously reported showing that large nonliving objects (9 mm in length) can be levitated between two square (5.0 × 5.0 × 2.5 cm) or disc (4.8 cm in diameter, 2.5 cm thick) permanent Neodymium (NdFeB) magnets larger than in microfluidic setups [37]. This technique has been then adapted to levitate millimeter-sized objects including living cell-laden beads and hydrogel units [38]. However, these configurations require the assembly of 2 NdFeB magnet blocks that constantly exert opposing forces on the system that hinders the technological translation of these setups for long-term usage. Furthermore, opposing magnets constrains the physical boundaries of the setup, limiting the access to the media for proper cell manipulation.

Recently, a ring magnet-based magnetic levitation configuration has been demonstrated for the density-based characterization of nonliving objects [39]. This configuration is composed of a single ring magnet and a glass tube of a paramagnetic solution, that are placed coaxially to each other, to provide better visualization and manipulation than that of the two-magnet configurations. Further, levitation systems composed of a pair of ring magnets with the same-poles facing have been proposed to engineer a linear, axially symmetric magnetic field for levitation and density-based analysis of nonliving and living objects [40–42]. Another magnetic installation containing a glass cuvette placed on an axial hole of two upright ring-shaped neodymium magnets with like poles facing each other was used for magnetic levitation of tissue spheroids in a paramagnetic medium [43]. Although these system designs allowed for a satisfying visualization of cell constructs, long-term culture of large living structures in a small culture volume, and performing routine cell culture operations such as media refreshment and recovery of samples, especially for mechanically unstable structures require additional considerations and remain untested.

Here, we showed the applicability of a one-step single ring magnet-based magnetic levitation design in formation and culture of 3D living structures. The system was shown to enable living cells to create large self-assembled 3D structures by preserving the cell viability and to allow mass transfers required for maintenance of the cell culture and combining biological units. We reported that the technique could be adapted for culture of several cell types and allowed transfer into intra-matrix culture. To the best of our knowledge, this is the first attempt to adapt a ring magnet-based magnetic levitation system for biofabrication of biological units and combining them.

## Materials and methods

### Design of magnetic levitation system

Magnetic levitation system is composed of a ring high grade (N52) neodymium (NdFeB) magnet (1” od × 5/16” id × 1/4” thick, K&J Magnetics) and a cell culture tube positioned in hole of the magnet (Figure 1). The bottom of the cell culture chamber is attached to the hole of the ring magnet with glue pads or the chamber is fixed on the magnet with a scaled photoreactive resin piece (Clear v2 FLGPCL02) printed using 3D printer (Formlabs Form 2). In the system, gadolinium (Gd^3+^) in the medium creates a difference (Δ_χ_ = *X*_c_ − *X*_m_) between magnetic susceptibility of the medium (*X*_m_) and cells’ (*X*_c_) to provide the levitation of cells where the magnetic force (*F*_mag_) acting on cells and the force of gravity (*F*_*g*_) balance each other. The magnetic forces directed to the centerline in the x-direction enables the cells to focus on the center for cellular aggregation.

**Figure 1.**
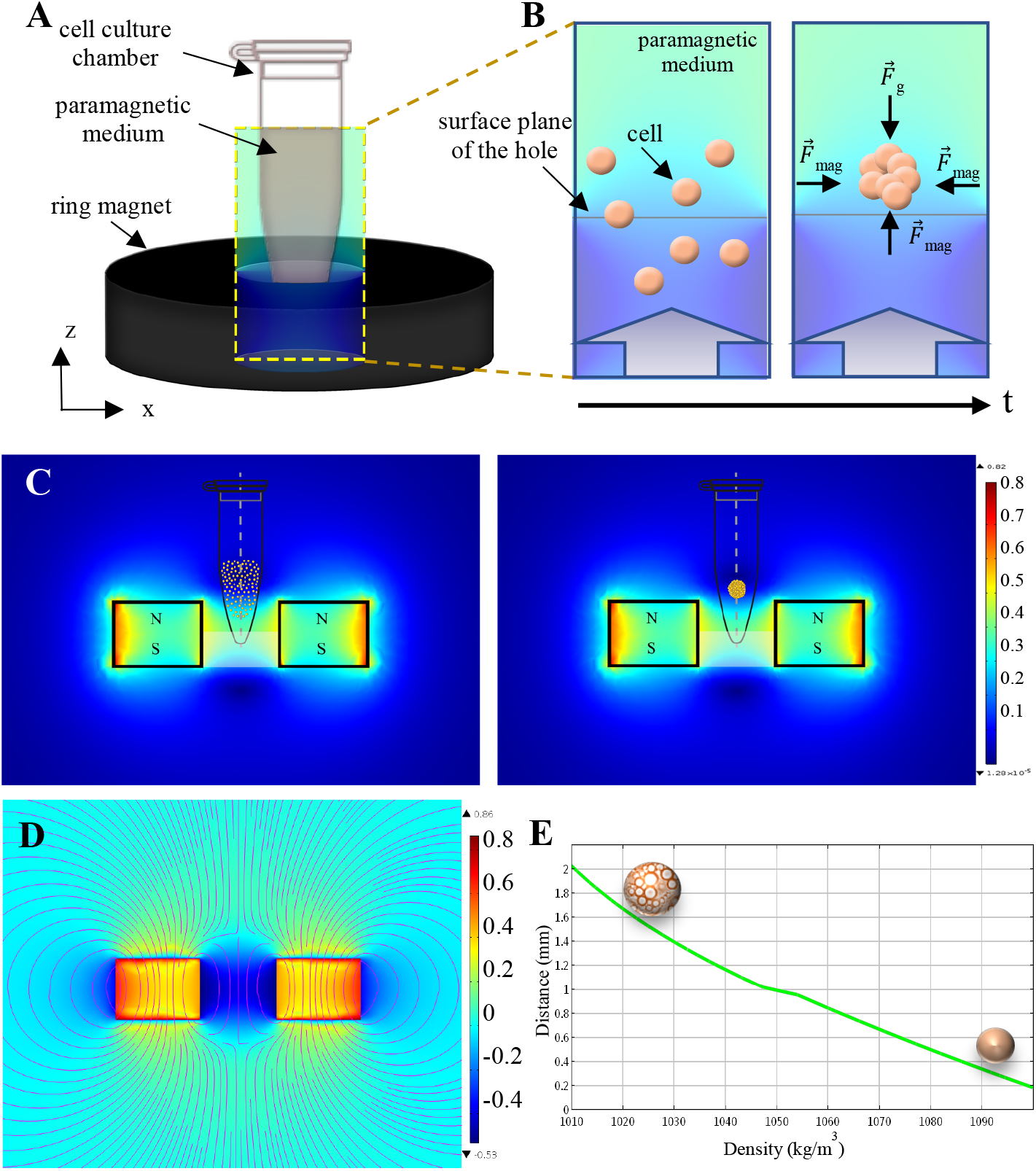
Magnetic force guided levitation and self-assembly. (A) Illustration of magnetic levitation system. Cell culture chamber is positioned on the ring magnet with bottom of the chamber attached to hole of the magnet. (B) Schematic representation of the cellular aggregation. The block arrows in the illustration represent upward magnetic induction. (C) Cellular aggregation represented on the simulation of magnetic flux density norm around the ring magnet. (D) Simulation of z component (B_z_) of magnetic flux density around the ring magnet via Finite Element Methodology. Total magnetic induction (B_z_+B_x_) is presented as streamlines. (E) Modeled relationship between the cell density and levitation heights for 200 mM concentration of Gd^3+^ based on the computational model. Level of the magnet surface is considered as z = 0. Density of cells as a function of their lipid content determines levitation height, and while less dense adipocytes are positioned at a higher level denser cells are positioned lower.

### Magnetic levitation of polymeric beads

Polymer beads with densities of 1.02 g/mL (size: 10–20 µm) and 1.09 g/mL (size: 20–27 µm) (Cospheric LLC., ABD), were suspended in the cell culture medium containing 0, 100 and 200 mM Gd^3+^ (Gadavist®, Bayer). Polymer bead suspension was loaded to a micro-capillary channel (1 mm × 1 mm square cross-section, 50-mm length, Vitrocom) and the channel was positioned on surface of the ring magnet by passing it over the hole of the magnet. That is to say the surface plane of the ring magnet serves as a ground for the levitation process. Movement of the beads in the magnetic field gradient was visualized under a stereo microscope (Soif Optical Instruments).

### Cell culture

D1 ORL UVA (bone marrow stem cell line, American Type Culture Collection (ATCC)) and MDA-MB-231 (breast cancer cell line, ATCC) cells were cultured in DMEM (Gibco) supplemented with 10% fetal bovine serum (FBS) and 1% penicillin/streptomycin. 7F2 (mouse osteoblasts, ATCC) were cultured in alpha modified essential medium (αMEM) supplemented with 10% FBS and 1% penicillin/streptomycin. The cells were grown in a humidified 37°C incubator with 5% CO_2_. The growth medium was refreshed every other day and the cells were passaged every four to six days. For adipogenic induction, 7F2 cells were exposed to induction medium composed of 5 µg/mL insulin, 10 nM dexamethasone and 50 mM indomethacin for 7 days. The induction medium was refreshed every other day. The cells were observed under an inverted microscope (Olympus IX-83).

### Levitation of living cells

The cells were detached with 0.25% trypsin-EDTA when the culture reached near confluency. Following centrifugation and removal of the supernatant, the cells were resuspended to 10^6^ cells/mL in the culture medium with various Gd^3+^ concentrations (50 mM, 100 mM, 150 mM, and 200 mM). 200 µL of cell suspension was loaded into the cell culture tube unless otherwise noted, and the culture tube was placed in the hole of the ring magnet. The cells were levitated in the magnetic levitation system for 24 hrs and imaged by a mobile phone equipped with a 15X micro focal length lens (Baseus) for short distance focusing. Horizontal diameter, vertical diameter, area and perimeter of the self-assembled clusters were measured with the ImageJ Fiji software.

### Visualization of trapping region for cellular cluster in the magnetic levitation system

D1 ORL UVA cells were resuspended to 10^6^ cells/mL in the culture medium with 200 mM Gd^3+^ and 100 µL of cell suspension was loaded into the cell culture chamber. Self-assembled cellular cluster after 48 hrs of levitation was moved upward and downward with the culture chamber to visualize cell trapping region in the vertical plane. For visualization of trapping region in the horizontal plane, the cell culture chamber was positioned horizontally on the ring magnet and moved parallel to the magnet surface until it passed the region where the movement of the cellular cluster was restricted. The motion of the cellular cluster was recorded by a mobile phone equipped with a 15X micro focal length lens.

### Modification of the medium and magnetic field

Ficoll® PM 400 (Sigma-Aldrich) was added to the culture medium to adjust the density of the medium to 1.02 and 1.04 g/mL. D1 ORL UVA cells (10^6^ cells/mL) were suspended in the denser culture media with 0 or 100 mM Gd^3+^. Levitation of cells in 200 µL were observed after 24 hrs of culture.

In order to further increase the magnetic susceptibility of the medium and thus the magnetic force on cells, levitation culture of D1 ORL UVA cells (10^6^ cells/mL) was performed with increasing concentrations of Gd^3+^; 0, 200, 350 and 500 mM. Levitation and aggregation of cells were observed within 5 hrs.

Two magnets have been attached with their opposite poles facing to strengthen the magnetic field in the levitation system. D1 ORL UVA cells were suspended in paramagnetic medium (150 or 200 mM Gd^3+^) at a concentration of 10^6^ cells/mL and levitated on holes of 1 ring magnet or 2 ring magnets whose opposite poles attached to each other. 200 µL of suspension was placed on levitation systems and cultures were observed after 2, 24 and 48 hrs.

### Transfer of cellular cluster and culture medium in magnetic levitation system

D1 ORL UVA cells were suspended in paramagnetic medium at a final concentration of 10^6^ cells/mL and 200 µL of cell suspension was cultured in magnetic levitation system for 48 h. To refresh the medium, old medium was removed with a pipette and fresh paramagnetic medium was slowly added to the culture.

To show the transfer of the resultant 3D culture, self-assembled compact clusters were harvested from the levitation culture without dispersion with a 1000 µL pipette tip. Harvested clusters were transferred to another levitation culture without dispersion with a 1000 µL pipette tip. All of the operations were recorded by a mobile phone equipped with a 15X micro focal length lens.

For culture maintenance of cellular cluster formed with magnetic levitation system in culture dish, D1 ORL UVA cells (10^6^ cells/mL) were levitated in 200 mM Gd^3+^ containing paramagnetic medium for 48 h and transferred to a culture dish with Gd-free medium. The culture was maintained for 24 h for observation.

### Live/Dead Assay

D1 ORL UVA cells were suspended in 200 mM Gd^3+^ containing paramagnetic medium and assembled in the magnetic levitation system. The levitation culture was maintained for 48 hrs before cell viability test. For viability test of the adipogenesis induced 3D structures, adipogenesis induced 7F2 cell were assembled in the magnetic levitation system for 24 hrs, then transferred to a culture plate and cultured for another 24 hrs. The viability of cells was assessed using live/dead assay (calcein-AM/propidium iodide, Sigma Aldrich) according to the manufacturer’s protocol. The cells were stained for 15 min at 37°C. Images were acquired using a fluorescence microscope (Olympus IX-83). The cells were both investigated as 3D cluster form and as dissociated single cells.

### Co-levitation culture

D1 ORL UVA cells were assembled and maintained during levitation by ring magnet-based magnetic levitation system for 48 hrs. The self-assembled spheres were transferred one by one to the ring magnet-based magnetic levitation system to a medium containing 200 mM Gd^3+^ in magnetic levitation system using a micropipette for co-levitation culture. Transfer of the clusters into the co-levitation culture was recorded by a mobile phone. For the co-levitation of 3D clusters consisting of lipid accumulated 7F2 cells, adipogenesis-induced cells were levitated in Gd^3+^-containing media at increasing concentrations (100, 150 and 200 mM) for 24 hrs, and the clusters were co-levitated in the same Gd-content medium in duplicate. Following a 24-hr co-levitation cultures, the 3D structures were transferred to the cell culture petri dish and observed under an inverted microscope (Olympus IX-83). Merged areas of each pair of spheres (%) were measured with ImageJ Fiji software by thresholding, followed by, shape completion and particle analysis.

### Embedding the 3D structures within the gel matrix

D1 ORL UVA cells were suspended in 200 mM Gd^3+^ containing medium and assembled in the magnetic levitation system for 24 hrs. At the end of the 24 hrs of the culture, the medium was aspirated until only 20 µL remained in the culture dish for maintenance of the levitation. Matrigel (BD Biosciences) with five times the volume of the remaining medium, was slowly added to the culture at +4°C and the culture was kept in a humidified 37°C incubator for 3 hrs. After the Matrigel polymerized, the 3D cell structure in the gel was transferred to a culture plate with the help of a pipette tip, which was cut on one side and turned into a micro spoon. Embedding the 3D structures within Matrigel and transfer of the clusters in Matrigel were recorded by a mobile phone. The medium was added onto the 3D structure embedded in Matrigel in culture dish and cultured for 4 days.

### Statistical analysis

All experiments were repeated at least three times. Results are reported as mean ± standard deviation (SD). Statistical analyses were performed using Student’s t-test (two-tail) or two-way analysis of variance (ANOVA) with Sidak post hoc correction, with GraphPad Prism version 6.0 (GraphPad Software). A p-value of < 5% was considered significant.

## Results

### Self-assembly of living cells in ring magnet-based magnetic levitation

A magnetic levitation system composed of a NdFeB (grade N52) ring magnet and a cell culture chamber was designed for levitation and self-assembly of cells (Figure 1(A)-(D)). First, we demonstrated that the system enabled levitation for the density range of the cells, 1.02-1.9 g / mL, and that the cells were localized at 0.3-1.7 mm distance from the magnet surface that was inversely proportional to their density based on the computational simulation (Figure 1(E)). In order to demonstrate the applicability of the ring magnet system for the levitation of living cells, polymeric beads with a density of 1.02 and 1.09 g/mL, representing the density of less dense and dense cells [13, 14], respectively were suspended in paramagnetic solution containing Gd^3+^ (100 and 200 mM), and their movements on the ring magnet were monitored (Supplementary Figure 1 and Supplementary Video 1-6). Polymeric beads with a density of 1.02 g/mL were levitated in paramagnetic media containing both 100 and 200 mM Gd^3+^, while denser particles (1.09 g/mL) were levitated in the medium containing 200 mM Gd^3+^, as they showed sedimentation at 100 mM Gd^3+^ concentration. After demonstrating that the system was able to provide levitation of particles with a density close to that of living cells, D1 ORL UVA cells were suspended in medium with increased concentrations of Gd^3+^ (0, 50, 100 and 200 mM) and cultured in the levitation system for 24 hrs (Figure 2). In the control group without Gd^3+^ all of the cells settled without levitation. In the paramagnetic medium containing 50 mM Gd^3+^, no cellular aggregates were formed. During the first 2 hrs of culture, the beginning of the cell clustering process in the paramagnetic medium with 100 and 200 mM Gd^3+^ concentrations were observable with the naked eye as cloudy aggregation of cells, and after 24 hrs, compact 3D structures were formed in these groups. While the majority of cells suspended in 100 mM could not be levitated in the system and sedimented to the bottom, cells suspended in 200 mM formed large 3D self-assembled clusters with levitation (Figure 2A red circles). The average horizontal diameter of cell clusters formed in medium with 200 mM Gd^3+^ was 867.33 ± 94.93 µm and approximately 1.7 times its vertical diameter. Cross-sectional area and perimeter of these clusters were measured as 0.39 ± 0.05 mm^2^ and 3.52 ± 0.36 mm, respectively (Figure 2B).

**Figure 2.**
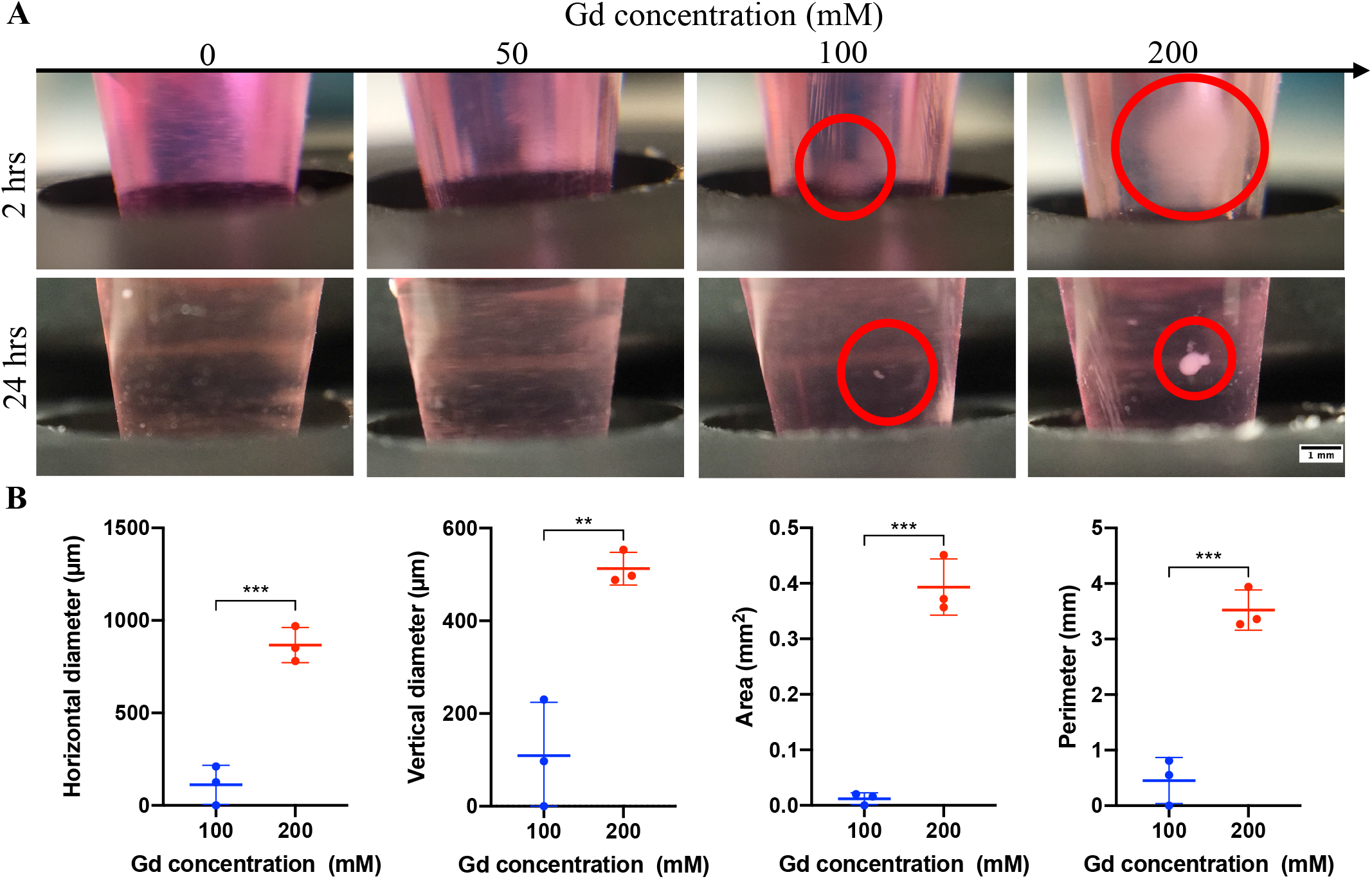
Self-assembly of D1 ORL UVA cells in ring magnet-based magnetic levitation system. (A) Micrographs of D1 ORL UVA cells cultured with ring magnet-based magnetic levitation (0, 50, 100 and 200 mM Gd^3+^, 10^6^ cells/mL, 100 µL) after 2 or 24 hrs of culture. Scale bar: 1 mm. (B) Size of the cellular clusters formed for 24 hrs with magnetic levitation (100 and 200 mM Gd^3+^, 10^6^ cells/mL, 100 µL); horizontal diameter, vertical diameter, area and perimeter. Data are plotted as mean of replicates with error bars (±SD) and statistical significance was determined by Student’s t-test (two-tail). **: p < 0.01; ***: p < 0.001.

Secondly, magnetic levitation culture was performed by manipulating the culture medium properties and the magnetic field. In order to reduce the gravitational force acting on the cells and therefore the magnetic susceptibility required to provide levitation, the density of the medium was increased by adding Ficoll to the medium, and D1 ORL UVA cells were levitated in these denser media. (Supplementary Figure 2). When the density of the culture medium was increased to 1.02 g/mL, the medium containing 100 mM Gd^3+^ concentration levitated most cells, unlike 1 g/mL medium. Measured horizontal diameter, vertical diameter, area and perimeter of cellular structures formed in medium with 1.02 g/mL density were 1005.33 ± 123.29 µm, 712 ± 54.03 µm, 0.70 ± 0.13 mm^2^ and 4.15 ± 1.09 mm, respectively. Moreover, horizontal diameter (p = 0.73), vertical diameter (p = 0.67), area (p = 0.24) and perimeter (p = 0.82) of cellular structures formed in medium with 1.02 g/mL density were statistically similar to structures observed with 1 g/mL medium with 200 mM Gd^3+^ concentration. Further increase in the medium density to 1.04 g/mL did not result in observable cluster formation.

To test whether rising the magnetic susceptibility of the medium increased the formation rate of cell clusters, we applied 350 and 500 mM Gd^3+^ concentrations, however no compact 3D structure was formed in any group within 5-hr levitation as in the media containing 200 mM Gd^3+^. (Supplementary Figure 3). Finally, we tested whether changing the strength of magnetic field by increasing lateral magnet area two-fold would affect the size of D1 ORL UVA cell clusters (Supplementary Figure 4(A)-(E)). However, biofabricated forms did not have a significant size difference in horizontal diameter (p = 0.62 and 0.74, respectively), vertical diameter (p = 0.50 and 0.56), area (p = 0.26 and 0.22) and perimeter (p = 0.99 and p = 0.57) in the medium containing 150 mM and 200 mM Gd^3+^. Computational simulation for magnet thickness implied that the system achieved a tighter focusing of cells with increased magnet thickness (Supplementary Figure 4(F)).

### Mass manipulation in 3D culture with ring magnet-based magnetic levitation

We next investigated the suitability of ring magnet-based magnetic levitation setup for mass manipulations in cell culture. In order to visualize cell focusing region in ring magnet-based magnetic levitation in the vertical plane, D1 ORL UVA cluster assembled with magnetic levitation was moved vertically with the culture chamber (Figure 3(A) and Supplementary Video 7). Equilibrium position was robustly kept by the cell cluster during the movement of the system in both directions. Next, to observe cell focusing region on the horizontal plane, D1 ORL UVA cluster formed by magnetic levitation culture was moved from the center of the magnet to the outside, parallel to the surface of the ring magnet with the culture chamber (Figure 3(B) and Supplementary Video 8). When the cellular structure reached the boundary of the area above the hole of the magnet, it was moved back towards the center of the magnet due to the high magnetic field on the magnet surface.

**Figure 3.**
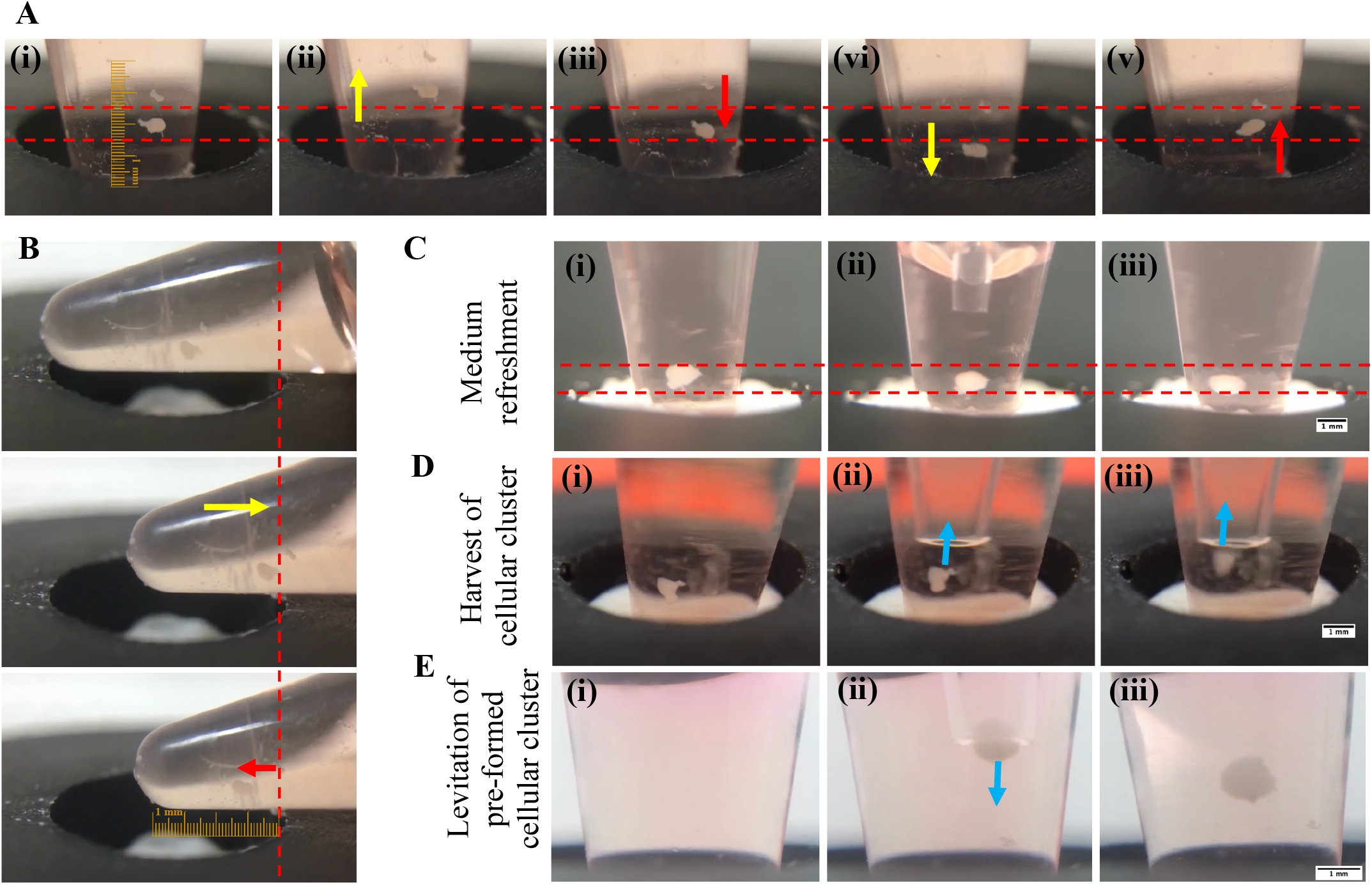
Mass manipulation in 3D culture with ring magnet-based magnetic levitation. (A) Trapping region of self-assembled D1 ORL UVA cluster (200 mM Gd^3+^, 10^6^ cells/mL, 100 µL) in the magnetic levitation system; (A) in the vertical plane, (B) in the horizontal plane. When the cellular cluster at equilibrium position (i) was moved upward with the culture chamber (ii), the cluster fell down into the equilibrium position (iii). When the cellular cluster was moved downward with the culture chamber (iv), the cluster rose back to its equilibrium position (v). Between the red dashed lines indicate the region in which the cellular cluster tends to be balanced in figure A. The red dashed line indicates the limit of the region in which the cellular cluster remains in the horizontal plane in figure B. Yellow arrows show the direction which the cellular cluster is moved with the culture chamber as an external force, and the red arrows show the direction which the cellular cluster inherently moves. (C) Refreshing culture medium of 3D cellular cluster formed in the magnetic levitation system (200 mM Gd^3+^, 10^6^ cells/mL, 200 µL) at the equilibrium position (i); removal of old medium (ii) and addition of fresh medium (iii). (D) Harvest of a 3D cellular cluster formed in the magnetic levitation system at the equilibrium position (i) by gently aspirating it with a pipette (ii, iii). (D) Transfer of 3D cellular cluster formed in the magnetic levitation system into another magnetic levitation device with a pipette. Scale bars: 1 mm.

Applicability of the medium refreshment, which is an essential factor for long term maintenance, was tested during levitation cell cultures that contained self-assembled and 48 hrs cultured D1 ORL UVA cells (Figure 3(C) and Supplementary Video 9). The ring magnet-based magnetic levitation system was found to be suitable for removing and replacing up to 80% of 200 µL total media volume with fresh medium, without causing the cellular cluster to settle. Gentle transfer of the medium using a micropipette ensured that the 3D structure in the system was not damaged. We followed this test of liquid phase transfer by viable cell cluster transfer. 3D structures formed of 2 × 10^5^ D1 ORL UVA cells were also gently collected from the levitation culture without being disturbed using a 1000 µL pipette tip (Figure 3(D) and Supplementary video 10), and the clusters that were harvested from a levitation culture were found to be transferred to another levitation culture without any disturbance (Figure 3(E) and Supplementary video 11).

### Long term levitation culture

Effects of levitation culture on the health of cells was tested by transferring 3D cellular spheres formed during 24 hrs of levitation culture to a standard culture dish (Figure 4(A) and Supplementary Figure 5). We observed that the cells spread adherently at the edges of the 3D cellular cluster to ∼43% of the cluster’s diameter for the sphere with a diameter of about 713 ± 3 µm. In order to determine the viability of the 3D structures formed in the magnetic levitation system, we performed a live/dead assay to both an intact 3D cluster (Figure 4(B)) as well as to a dissociated form as a single-cell suspension (Figure 4(C)). Visual inspection of the live/dead fluorescence microscopy images showed that most of the cells were viable as apparent from the green calcein-AM signal in both 3D form and the single cell suspension.

**Figure 4.**
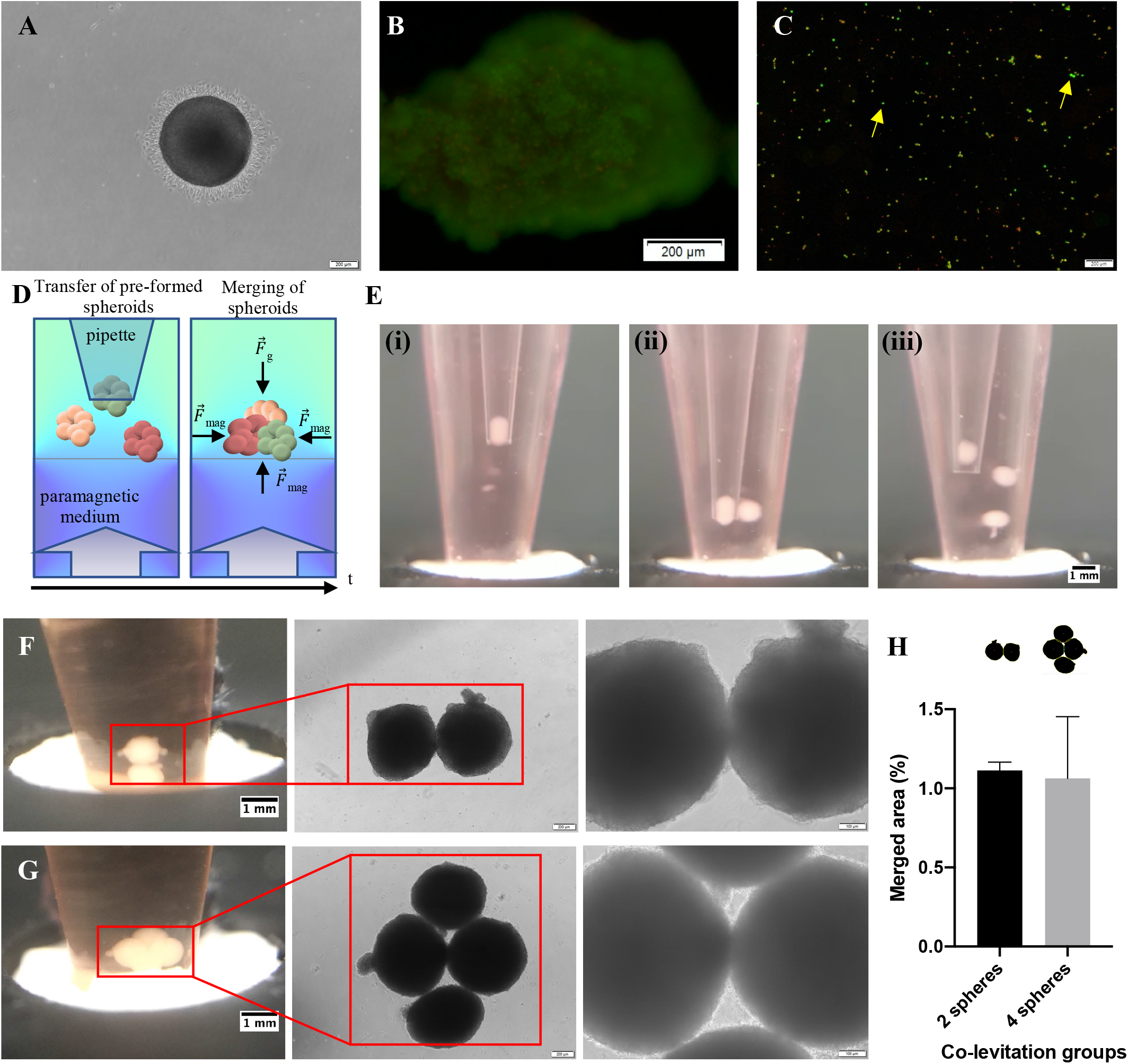
Post-operations on spheres formed by ring magnet-based levitation. (A) Micrograph of a self-assembled D1 ORL UVA 3D structure cultured with magnetic levitation (200 mM Gd^3+^, 10^6^ cells/mL, 200 µL) for 48 hrs and then cultured for 24 hrs in the 2D culture dish. Fluorescent microscopy images of D1 ORL UVA (B) 3D structures formed with magnetic levitation and (C) cells dissociated from the 3D structures. (live: green, dead: red). Cell viability was visualized by live-dead staining (Calcein/PI). Yellow arrows denote some of alive cells. Scale bar: 200 µm. (D) Schematic representation of the co-levitation of self-assembled cellular clusters. (E) One-by-one transfer of D1 ORL UVA cellular clusters that were individually self-assembled and cultured for 48 hrs in ring magnet-based magnetic levitation system to the medium containing 200 mM Gd^3+^ in magnetic levitation system for co-levitation culture. Co-levitation culture of preformed (F) two and (G) four D1 ORL UVA cellular clusters in medium containing 200 mM Gd^3+^ in the magnetic levitation system for 24 hrs. Scale bars: 1 mm for culture chamber images, 200 and 100 µm for middle and right images, respectively, in F and G. (H) Merged area of spheres (%) co-levitated in a medium containing 200 mM Gd^3+^ for 24 hrs. Data are plotted as mean of replicates with error bars (±SD) and evaluated using the unpaired Student’s t-test.

The potential of the ring magnet-based magnetic levitation system to biofabricate complex structures consisting of 3D living units was demonstrated by successful co-levitation of homocellular spheroids that were levitation cultured and transferred from a prior system (Figure 4(D)-(G)). The 3D spheres formed as a result of magnetic levitation of D1 ORL UVA cells were gently transferred into the medium containing 200 mM Gd^3+^ in the levitation system (Figure 4(E)). Co-levitation cultures formed by transferring two or four of them to the device (Supplementary Video 12 and 13) were maintained for another 24 hrs to allow cell–cell attachment between spheroids. We observed that the cellular spheres were fused after 24 hrs of co-levitation and they were successfully transferred to a different culture dish without deterioration for a better display of the inter-cluster contact zones in 3D structures (Supplementary Figure 6). A 24 hrs co-levitation resulted in 1.07 ± 0.35% merging of the spheres in area and no statistical difference was observed between percentage fusion in bilateral and quadruple co-levitation cultures (p=0.87) (Figure 4(H) and Supplementary Figure 7).

### Magnetically guided self-assembly of cells with different single cell densities

Since one of the features determining the final position of the cells in the magnetic levitation principle is the inherent single cell densities, the levitation-based 3D culture protocol of low-density cells in the system was defined using adipocytes with low density due to cellular lipid accumulation [14]. Adipogenesis of 7F2 cells were induced for 7 days to obtain lipid accumulated cells (Supplementary Figure 8). Following the observation of lipid accumulation, the cells were suspended in the paramagnetic medium containing increasing Gd^3+^ concentrations (0, 100, 150 and 200 mM) and levitation cultured over 24 hrs in the ring magnet-based magnetic levitation system (Figure 5(A)). We observed that the cells started to accumulate on the magnet towards the center in all paramagnetic medium containing Gd^3+^ between 100 and 200 mM at the 2^nd^ hr of the culture, and these stably levitated cells formed 3D structures at the 24^th^ hr of the culture. The spheres formed in 100 mM Gd^3+^ containing medium were 1.95 and 2.95 times larger in area (∼ 5.8 mm^2^), and 1.51 and 1.58 times larger in perimeter (∼ 10.72 mm) than those formed in the medium containing 150 and 200 mM Gd^3+^, respectively (Figure 5(B)). There was no statistically significant difference between the areas (p=0.06) and perimeters (p=0.78) of the clusters formed in the medium containing 150 and 200 mM Gd^3+^. When the cells were assembled in the medium containing 100 mM Gd^3+^, the shapes of the clusters were skewed in the direction of the vertical diameter rather than horizontal diameter compared to the clusters formed in paramagnetic medium containing higher Gd^3+^. Closer inspection of the graph showed that the vertical diameter of the cellular clusters formed in the medium containing 100 mM Gd^3+^ was 3917 ± 622.55 µm and it was 2.38 and 2.72 times more than the clusters formed at 150 mM and 200 mM Gd^3+^ concentrations, respectively.

**Figure 5.**
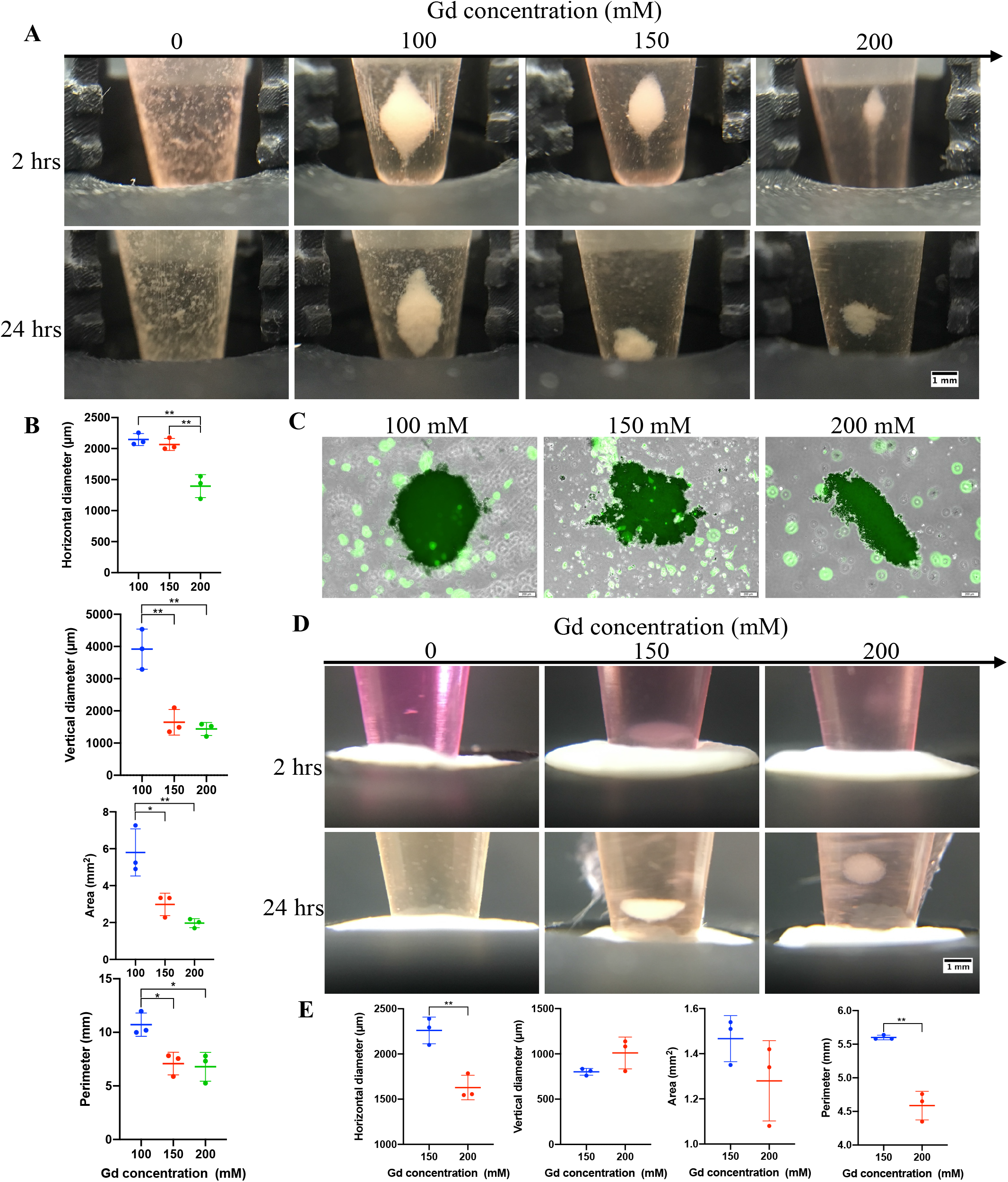
Levitation based 3D culture of different cell types. (A) Micrographs of adipogenesis induced 7F2 cells cultured with ring magnet-based magnetic levitation (0, 100, 150 and 200 mM Gd^3+^, 10^6^ cells/mL, 200 µL) after 2 or 24 hrs of culture. Each vertical unit on the 3D printed scaled piece: 1mm. Scale bar: 1 mm. (B) Size of the adipogenesis induced 7F2 cellular clusters formed for 24 hrs with magnetic levitation; horizontal diameter, vertical diameter, area and perimeter. (C) Fluorescent microscopy images of adipogenesis induced 7F2 3D structures formed with magnetic levitation. Cell viability was visualized by live staining (Calcein-AM). Scale bar: 200 µm. (D) Micrographs of MDA-MB-231 cells cultured with ring magnet-based magnetic levitation (0, 150 and 200 mM Gd^3+^, 10^6^ cells/mL, 200 µL) after 2 or 24 hrs of culture. Scale bar: 1 mm. (E) Size of the MDA-MB-231 cellular clusters formed for 24 hrs with magnetic levitation; horizontal diameter, vertical diameter, area and perimeter. Data are plotted as mean of replicates with error bars (±SD) and statistical significance was determined by Student’s t-test (two-tail). *: p < 0.05; **: p < 0.01.

In order to maintain culture of the adipogenesis induced 3D structures, which were formed as a result of 24-hr levitation, spheroids were transferred to a culture plate and cultured for another 24 hrs (Supplementary Figure 9). We observed that the transferred 3D structures were loose and many adipocytes dissociated from the edges of the 3D clusters in all paramagnetic medium groups after the transfer. While most of the cells separated from the main cluster were in suspended form, some lipid-containing cells were observed to spread over the culture surface. Testing the viability of cells at the end of the culture by live/dead staining showed that most cells in the 3D cluster were alive (Figure 5(C) and Supplementary Figure 10). We also carried out co-levitation of 3D adipogenesis-induced cell clusters formed separately in the same medium for 24 hrs (Supplementary Figure 11). Although the clusters appeared together with the assistance of magnetic force in the levitation system at the 24^th^ hr of levitation, we observed that there was still no fusion between the clusters when transferred to the culture vessel and the clusters were dispersed with the transfer.

The ring magnet-based magnetic levitation system was also tested for biofabrication of 3D structures of MDA-MB-231 breast cancer cells levitation cultured for 24 hrs in medium containing 150 and 200 mM Gd^3+^ (Figure 5(D)). In the 2^nd^ hr of the culture, cell clustering began with a nebulous appearance in the paramagnetic medium and levitated tight 3D clusters with an area of 1.37 ± 0.17 mm^2^ were observed at 24^th^ hr (Figure 5(E)). The horizontal diameter of the 3D structures formed in the media containing 150 mM Gd^3+^ was ∼39% higher and the perimeter was ∼22% higher than those formed in the medium containing 200 mM Gd^3+^.

### In-gel culture of self-assembled 3D structures

In order to demonstrate the transferability and the sustainability of self-assembled 3D structures created with magnetic levitation into an in-gel culture, D1 ORL UVA cells were levitation cultured for 24 hrs and at the end of the culture self-assembled 3D structures were embedded in Matrigel (Figure 6(A), (B) and Supplementary Video 14). After aspirating most of the levitation medium leaving enough to sustain the levitation of the 3D structure (∼20 µL) we slowly added Matrigel to the levitation system. We transferred the Matrigel in a slow rate to the point that the Matrigel volume was five times the volume of the remaining medium. Matrix was added at +4°C that keep it in liquid form and polymerization was achieved by temperature change. It was shown that the levitated cellular structures could be successfully trapped within the Matrigel without any observable deformation. We then transferred the cellular structure within the gel matrix to a separate culture (Figure 6(C)-(E) and Supplementary Video 15). On the 4^th^ day of the culture, we observed that the 3D cellular structure consisted of viable cells spreading in the gel matrix (Figure 6(F), Supplementary Figure 12).

**Figure 6.**
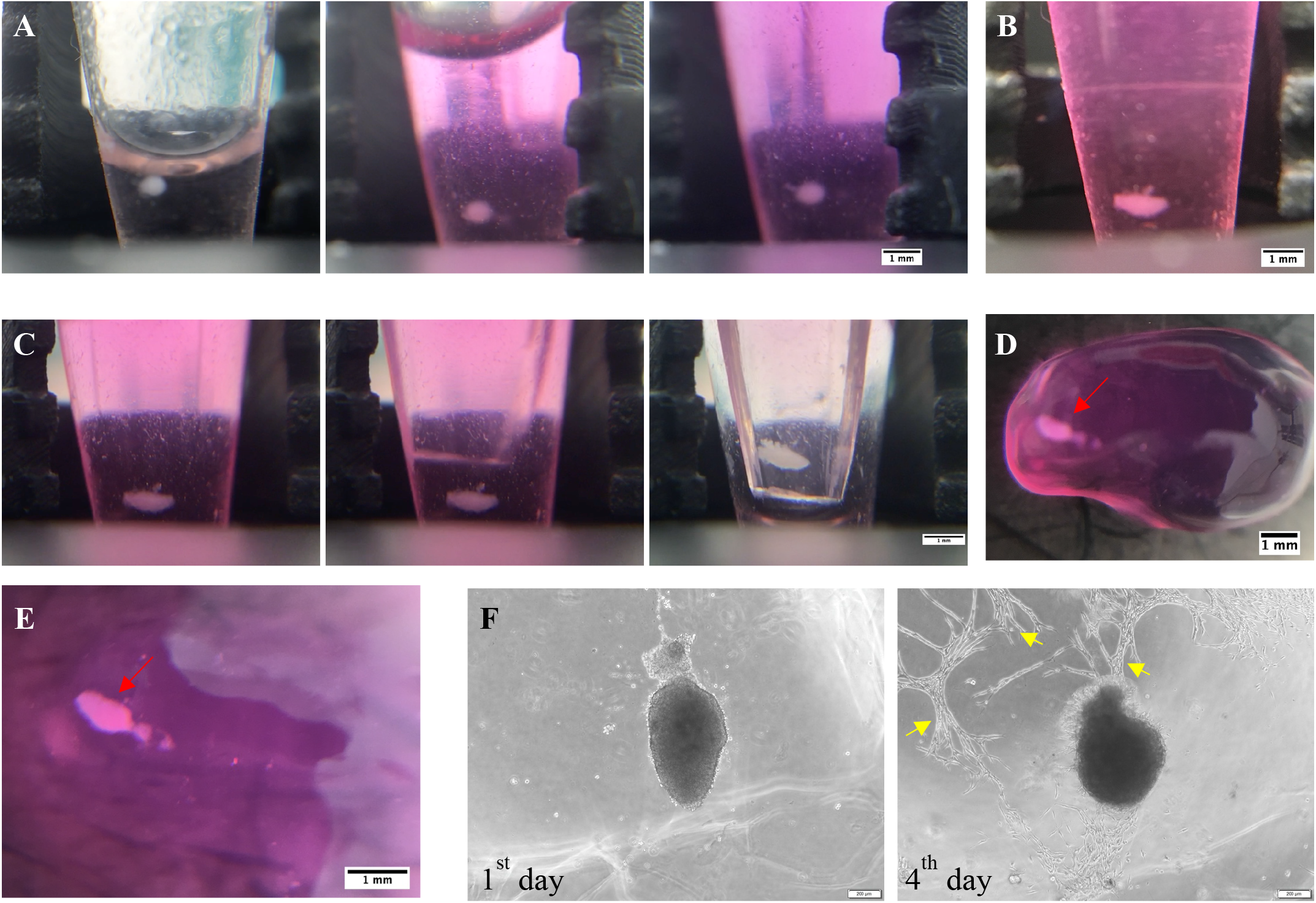
In-gel culture of self-assembled D1 ORL UVA 3D structures. (A) Embedding a 3D cellular structure assembled by magnetic levitation within Matrigel. (B) Cellular cluster in Matrigel at 3^rd^ hr of culture. (C) Harvest of gel-embedded 3D cluster using a pipette tip, which was cut into a micro-spoon. Matrigel-embedded 3D cluster that was transferred into a culture plate. (D) Before and (E) after medium addition on gel-embedded culture. Red arrows show the 3D clusters in the gel matrix. Each vertical unit on the 3D printed scaled piece: 1mm. Scale bar: 1 mm (F) Micrographs of the Matrigel-embedded 3D cluster after 1 and 4 days of culture. Yellow arrows indicate some of the cells spreading in the gel matrix. Scale bar: 200 µm.

## Discussion

Magnetic force-assisted cell manipulation provides a broadly applicable guidance tool in many fields such as biological or clinical research and tissue engineering. The availability of label-free protocols has recently led to a greater focus of research on these techniques due to both lowering required cost, time and labor, and enhancing compatibility of the technique for living cells. Some microcapillary based magnetic levitation systems, that were initially applied to detect and sort cells of interest according to their physical intrinsic properties [13, 14, 44–46], were later adapted for biofabrication [19, 20, 47]. While great progress has been made in the field, tissue engineering applications and biological testing protocols require manufacture of sizable living constructs to provide an adequate number of cells. Systems that allow cell culture applications on-site and offer low-cost applications with permanent magnets are essential to render the production easy to install and operate, and to enable on-site intervention to production process.

The standard diamagnetic levitation devices using capillary tubes (1 mm × 1 mm square cross-section) physically sandwiched between two block permanent magnets are able to create large cellular blocks (up to ∼2.68 cm in length) [19]. Although these elongated living structures created in such systems are advantageous in terms of efficient mass transfer between cluster and its surrounding, they are not mechanically resistant to transfer processes due to their low thickness (up to ∼280 µm) with low homogeneity towards the capillary ends, limiting their applications in bottom-up tissue engineering. Another magnetic levitation setup design was shown to increase the working volume by positioning larger block magnets (poles on 2 inch × 2 inch surfaces) further apart with a gap set to 2.5 cm [38]. This system allows remote 3D manipulation of millimeter-sized living objects. Such systems still contain opposing magnets occupying the top and bottom of the culture chamber to provide magnetic field gradient required for levitation, and this configuration limits operations on the culture during levitation process such as medium refreshment and transfer of the cellular structures. Ring magnet-based magnetic levitation system proposed here removes the upper physical barrier, hence providing an easy access to the levitating biological structures and to its surrounding medium. Furthermore, this setup eliminates the limit for the height of the cell culture chamber and thus enables levitation in great height culture containers. We showed that single step axial-circular magnetic levitation made addition and removing of liquid or solid phases straightforward without removing the culture chamber from the magnetic field owing to a large and open operational space on the culture container. The ability to be processed during levitation also provided the opportunity to fully embed the levitated structures in another phase such as a gel matrix. The sustainability of the culture within a gel matrix ensures that the system can be applied effectively in broad studies including drug response, cell movement and stromal effects. Mammalian cells exhibit different characteristics for densities depending on their type; e.g. ∼1.044 g/mL for breast cancer cells, ∼1.062 g/mL for lung cancer cells [13], ∼1.084 g/mL for bone marrow originated stem cells [14]. Here we have shown that ring magnet-based magnetic levitation system is able to levitate objects with a density ranging from 1.02-1.09 g/mL by levitation of particles with known density. Considering the variability of density depending on cellular condition such as type of cell, pathological conditions and differentiation [48], the wide range of applicability of the system has been demonstrated. As models representing levitation of cells with different densities, stem cells, breast cancer cells and adipocytes were self-assembled into 3D structures with preserving cell viability in our system. It was shown that tight and intact 3D cellular units were produced with bone marrow originated stem cells and breast cancer cells and the magnetic levitation system could provide the fusion of biological units composed of stem cells. However, 3D adipocyte clusters were mechanically too unstable for transfer and fusion operations. As the system relies on cell-cell interaction independent of an external mechanical support, the technique is not suitable for natural self-assembly of each cell type. These loose structures may be modified and strengthened by using binary cell mixtures including fibroblasts [49] and stem cells [20] to act as an adhesive that promotes intercellular interactions.

The process of creating 3D cellular structures by magnetic guidance involves first focusing the single cells homogeneously distributed in the suspension by magnetic force and then gaining a stable architecture of the structure with cell-cell interactions in the focusing region. 3D stable structure formation on the ring magnet-based magnetic levitation system presented herein took more than 10 hours regardless of cell type. In order to shorten this period into couple hours, the magnetic force applied on the cells was increased by modifying the paramagnetic media, however the formation process could not be accelerated. Previously a magnetic manipulation method has been described to print 3D cellular structures within 6 hours [50]. Unlike our system, this method enabled individual cells to focus on the culture surface rather than levitational assembly. In order to accelerate cellular aggregation in the ring magnet-based levitation system, the cell focusing process may be accelerated by the physical confinement of the cells in the region close to the low magnetic field (e.g. increasing magnet thickness) or by using binary cell mixtures as an adhesive.

Mini-tissue block fabrication shows great promise in formation of complex and large 3D anatomical structures. Tissue blocks, which create their own matrix and architecture, show a potential as building blocks for scale-up tissue fabrication. The absence of an external material allows biomaterial-based concerns such as material-induced toxicity and host inflammatory responses to be overcome [51–54]. However, while scaffolding provides void volume for passive diffusion of nutrients, gasses and wastes into the scaffolds to keep the cells alive [55], scaffold-free approaches lack this advantage. Our axial-circular magnetic levitation system may be equipped with a flow system that the bulk liquid phase is continuously refreshed to improve the diffusion between the sphere surface and the liquid phase without disturbing the levitation. In the case of increased spherical size and extended culture periods, the protocol should be tested for the health of cells in the central region prior to fabrication.

## Conclusions

In this study, we proposed an axial-circular magnetic levitation system to remotely manipulate living cells for biofabrication. We verified that the single ring magnet-based magnetic levitation system enabled fabrication of living building blocks and their fusion based on the cellular self-assembly as an alternative to commonly used magnetic levitation systems composed of a culture chamber between two block magnets. The proposed magnetic levitation configuration provides several advantages: (i) Levitation on a single ring magnet eliminates the limit on the height of the culture reservoir, making the system broadly compatible for different types of culture chambers like tubes and cuvettes, and allowing to form and maintain sizeable living structures. (ii) The system provides an open and large operational space allowing for easy and on-site intervention in cell culture such as medium refreshment and adding another structure without disturbing levitation and structures. (iii) Permanent magnets are common and electrical power independent, therefore the design enables straightforward, simple and low-cost installation and operation. (iv) Label-, scaffold- and nozzle-free nature of the protocol allows for manufacturing of living constructs that is rapid, natural-like and free from mechanical stress. The platform presented here may be improved by automation with a perfusion system for a continuous fabrication and by providing on-site monitoring at high magnification under microscope using mirrors. The system offers wide range applications including biofabrication of scale-up complex structures and of tissue models for drug testing and cancer research by operating in batch or continuous mode.

## Acknowledgments

Financial support by The Scientific and Technological Research Council of Turkey (119M755) is gratefully acknowledged.

## Figure captions

**Supplementary Figure 1.**
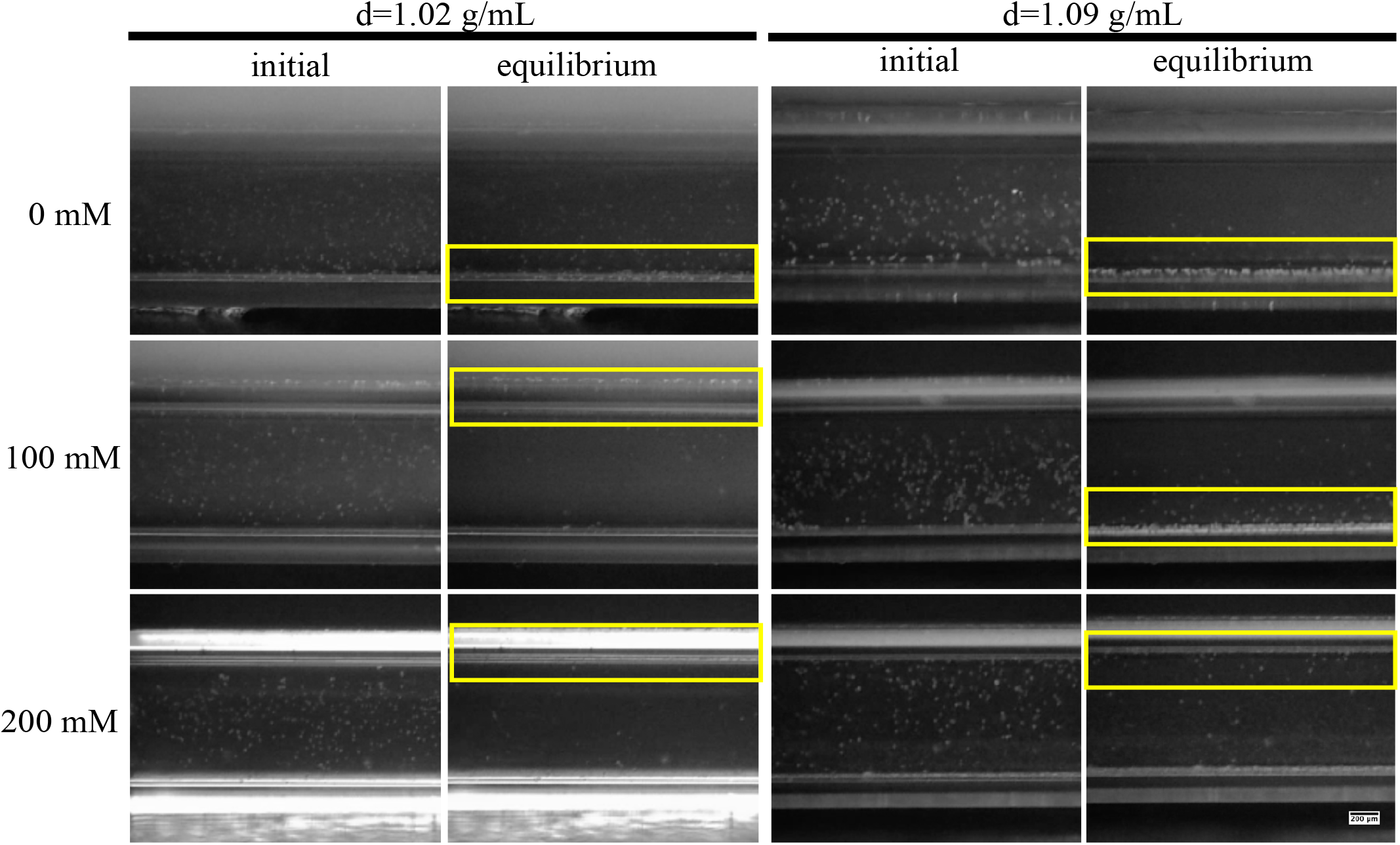
Micrographs of polymeric beads with density of 1.02 and 1.09 g/mL suspended in the culture medium (0, 100 and 200 mM Gd^3+^) on the hole of ring magnet. The first micrographs were recorded at the beginning of the process, and the second micrographs were recorded when the beads reached equilibrium position (within 7 min). Yellow rectangles show the region where the cells were collected. Scale bar: 200 µm.

**Supplementary Video 1**. Real-time monitoring of polymeric beads with a density of 1.02 g/mL suspended in the culture medium (0 mM Gd^3+^) on the hole of ring magnet.

**Supplementary Video 2**. Real-time monitoring of polymeric beads with a density of 1.02 g/mL suspended in the paramagnetic medium (100 mM Gd^3+^) on the hole of ring magnet.

**Supplementary Video 3**. Real-time monitoring of polymeric beads with a density of 1.02 g/mL suspended in the paramagnetic medium (200 mM Gd^3+^) on the hole of ring magnet.

**Supplementary Video 4**. Real-time monitoring of polymeric beads with a density of 1.09 g/mL suspended in the culture medium (0 mM Gd^3+^) on the hole of ring magnet.

**Supplementary Video 5**. Real-time monitoring of polymeric beads with a density of 1.09 g/mL suspended in the paramagnetic medium (100 mM Gd^3+^) on the hole of ring magnet.

**Supplementary Video 6**. Real-time monitoring of polymeric beads with a density of 1.09 g/mL suspended in the paramagnetic medium (200 mM Gd^3+^) on the hole of ring magnet.

**Supplementary Figure 2.**
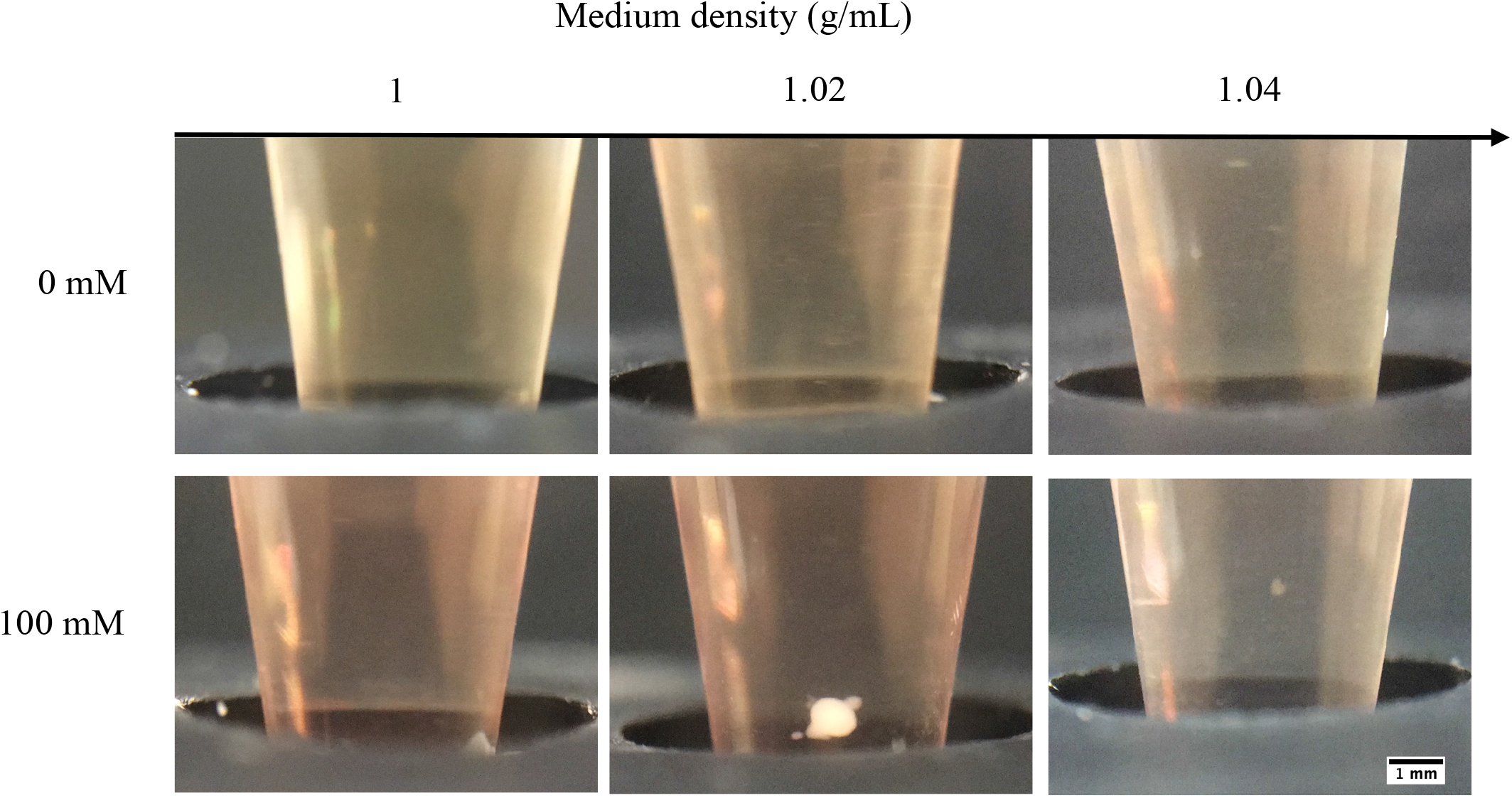
Micrographs of D1 ORL UVA cells levitated in culture medium with densities of 1 (without Ficoll), 1.02 and 1.04 g/mL with ring magnet-based magnetic levitation (0 and 100 Gd^3+^) after 24 hrs of culture. Scale bar: 1 mm.

**Supplementary Figure 3.**
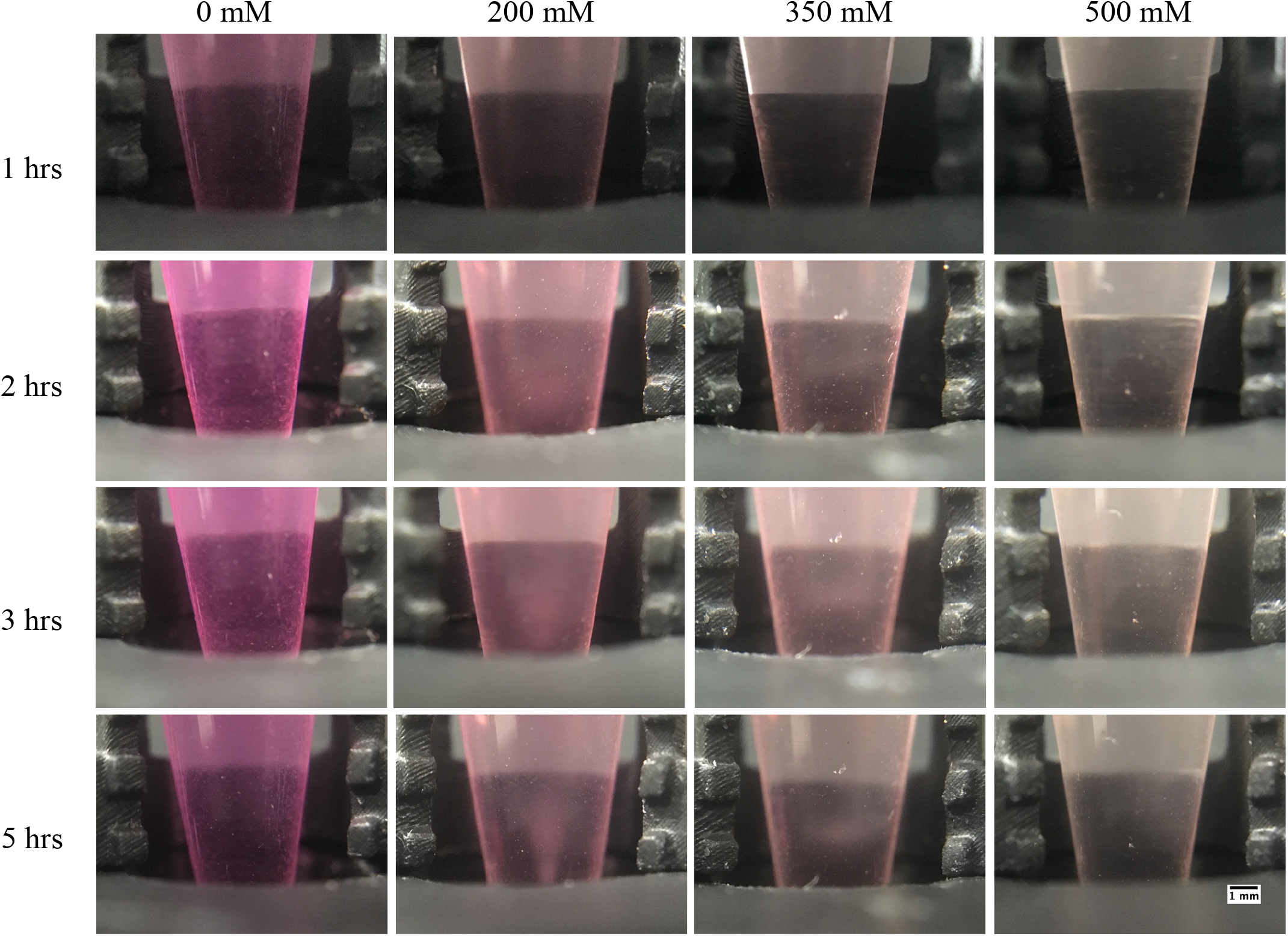
Micrographs of D1 ORL UVA cells cultured with increasing concentrations of Gd^+3^ (0, 200, 350 and 500 mM) in the ring magnet-based magnetic levitation platform within 5 hrs of culture. Each vertical unit on the 3D printed scaled piece: 1mm. Scale bar: 1 mm.

**Supplementary Figure 4.**
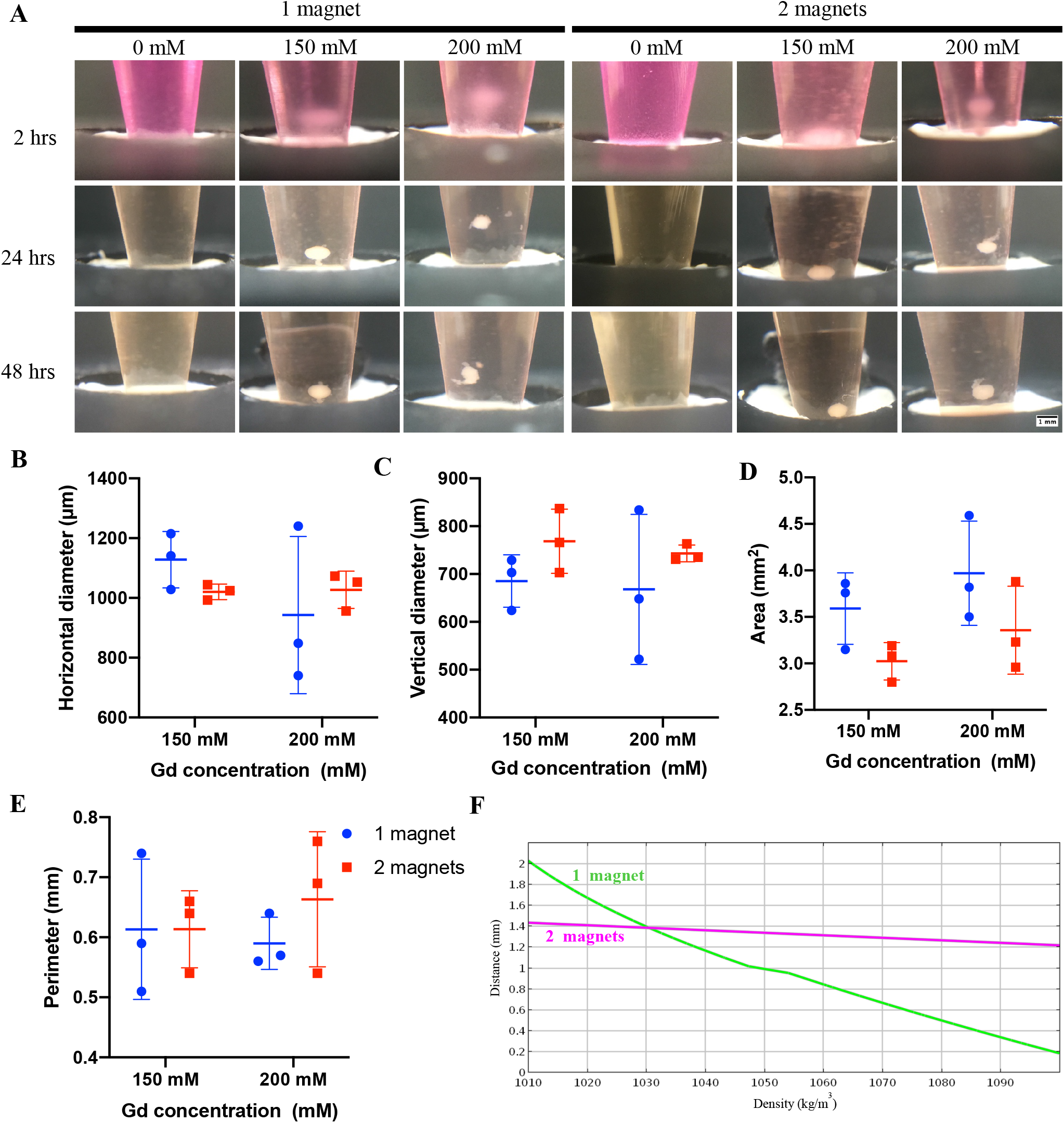
(A) Micrographs of D1 ORL UVA cells cultured in the ring magnet-based magnetic levitation platforms (0, 150, 200 mM Gd^+3^) composed of 1 ring magnet or 2 ring magnets whose opposite poles attached to each other after 2, 24 and 48 hrs of culture. Scale bar: 1 mm. (B-E) Size of the cellular clusters formed for 24 hrs in the ring magnet-based magnetic levitation platforms (150, 200 mM Gd^+3^) composed of 1 ring magnet or 2 ring magnets; horizontal diameter, vertical diameter, area and perimeter. Data are plotted as mean of replicates with error bars (±SD) and statistical significance was determined by two-way ANOVA with Sidak post hoc correction. (F) Modeled relationship between the cell density and levitation heights in the ring magnet-based magnetic levitation platforms (200 mM Gd^+3^) composed of 1 ring magnet or 2 ring magnets based on the computational model. Level of the magnet surface is considered as z = 0.

**Supplementary Video 7**. Movement of self-assembled D1 ORL UVA cluster in ring magnet-based magnetic levitation system following forced movement of the cellular cluster with the culture chamber in the vertical plane.

**Supplementary Video 8**. Movement of self-assembled D1 ORL UVA cluster in ring magnet-based magnetic levitation system following forced movement of the cellular cluster with the culture chamber in the horizontal plane.

**Supplementary Video 9**. Refreshing culture medium of self-assembled 3D cellular cluster in the ring magnet-based magnetic levitation system.

**Supplementary Video 10**. Harvest of a self-assembled 3D cellular cluster formed in the ring magnet-based magnetic levitation system.

**Supplementary Video 11**. Transfer of a self-assembled formed in the magnetic levitation system into another ring magnet-based magnetic levitation device.

**Supplementary Figure 5.**
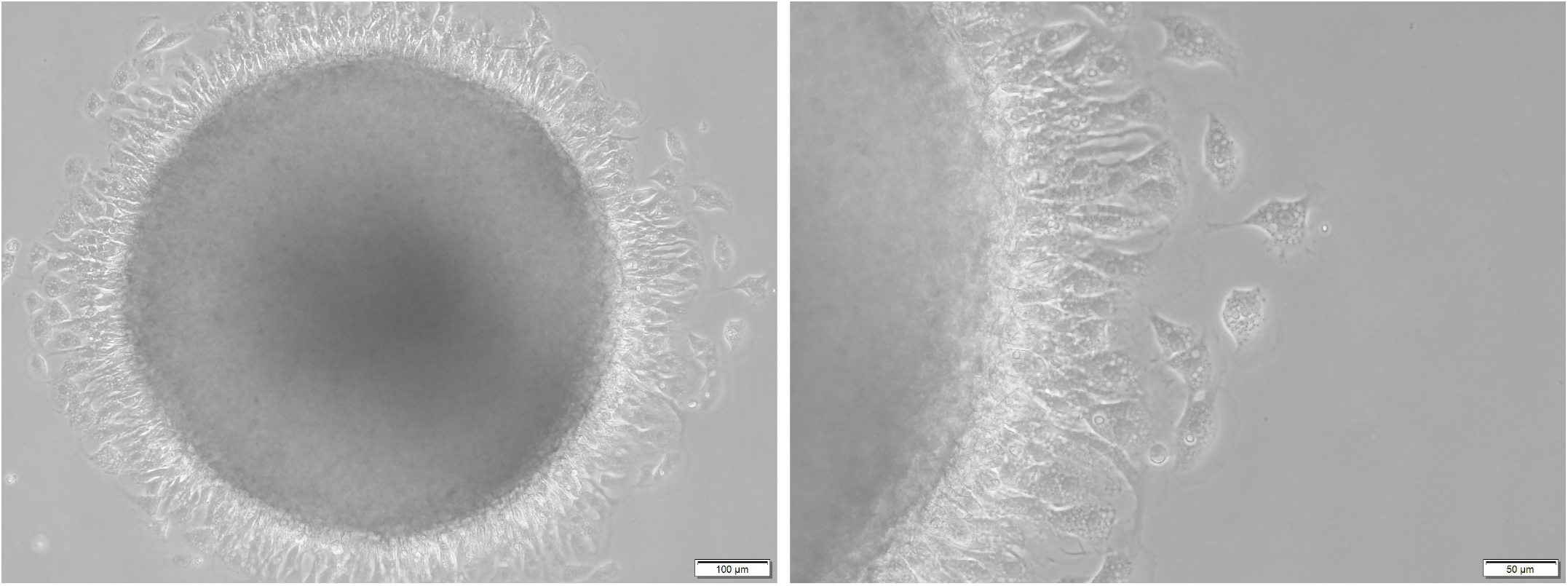
Micrograph of a self-assembled D1 ORL UVA 3D structure cultured with magnetic levitation (200 mM Gd^3+^, 10^6^ cells/mL, 200 µL) for 48 hrs and then cultured for 24 hrs in the culture dish. Scale bars: 100 and 50 µm, respectively.

**Supplementary video 12**. One-by-one transfer of two D1 ORL UVA cellular spheroids that were individually self-assembled and cultured for 48 hrs in ring magnet-based magnetic levitation system to the medium containing 200 mM Gd^3+^ in the magnetic levitation system for co-levitation culture.

**Supplementary video 13**. One-by-one transfer of four D1 ORL UVA cellular spheroids that were individually self-assembled and cultured for 48 hrs in ring magnet-based magnetic levitation system to a medium containing 200 mM Gd^3+^ in the magnetic levitation system for co-levitation culture.

**Supplementary Figure 6.**
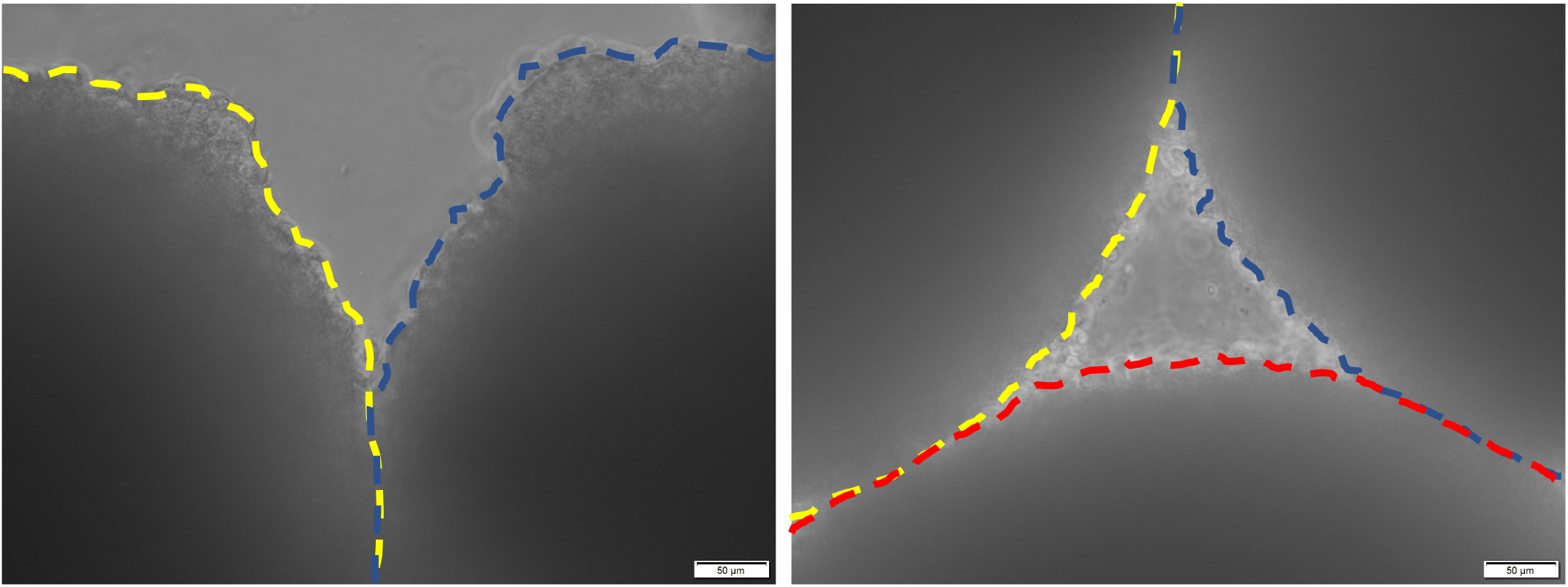
Zoomed-in view of the inter-cluster contact zones in 3D structures formed by 24-h co-levitation of two and four D1 ORL UVA cellular clusters in medium containing 200 mM Gd^3+^. Dashed lines indicate the boundaries of the spheroids. Scale bar: 50 µm.

**Supplementary Figure 7.**
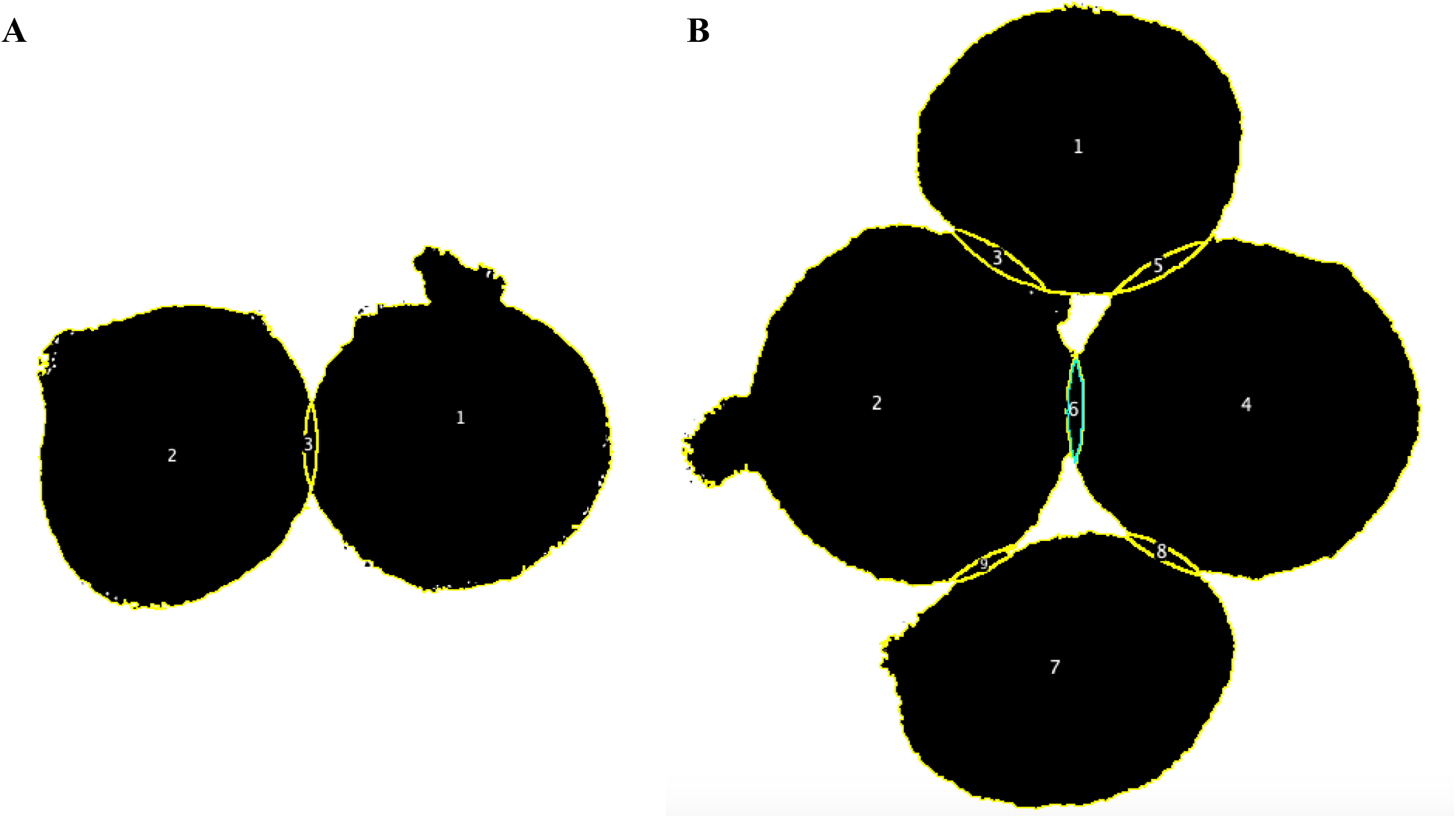
Images obtained for calculation of the merged areas (%) represented in figure 4 (H) by an image processing for co-levitation culture of preformed (A) two and (B) four cellular clusters.

**Supplementary Figure 8.**
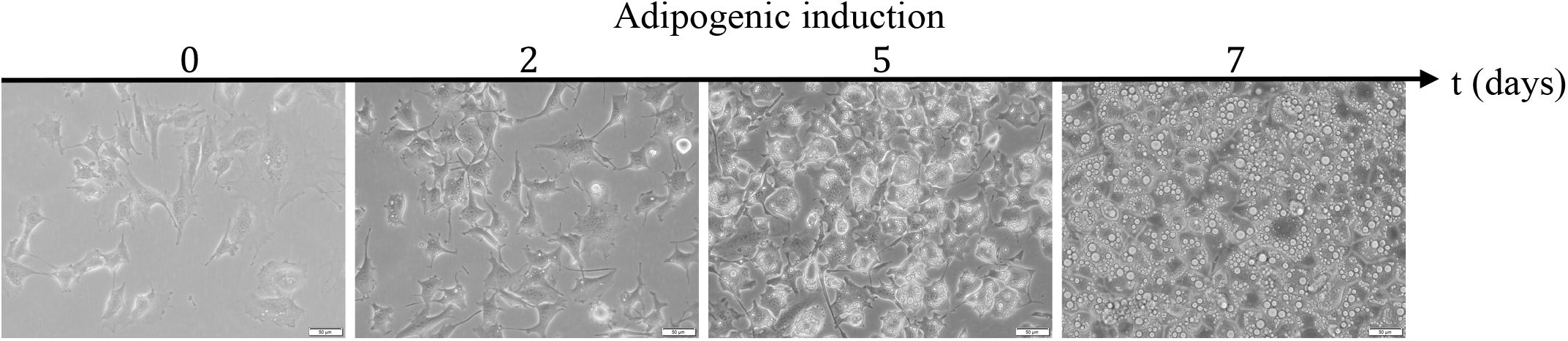
Micrographs of adipogenesis induced 7F2 cells over 7 days. Scale bar: 50 µm.

**Supplementary Figure 9.**
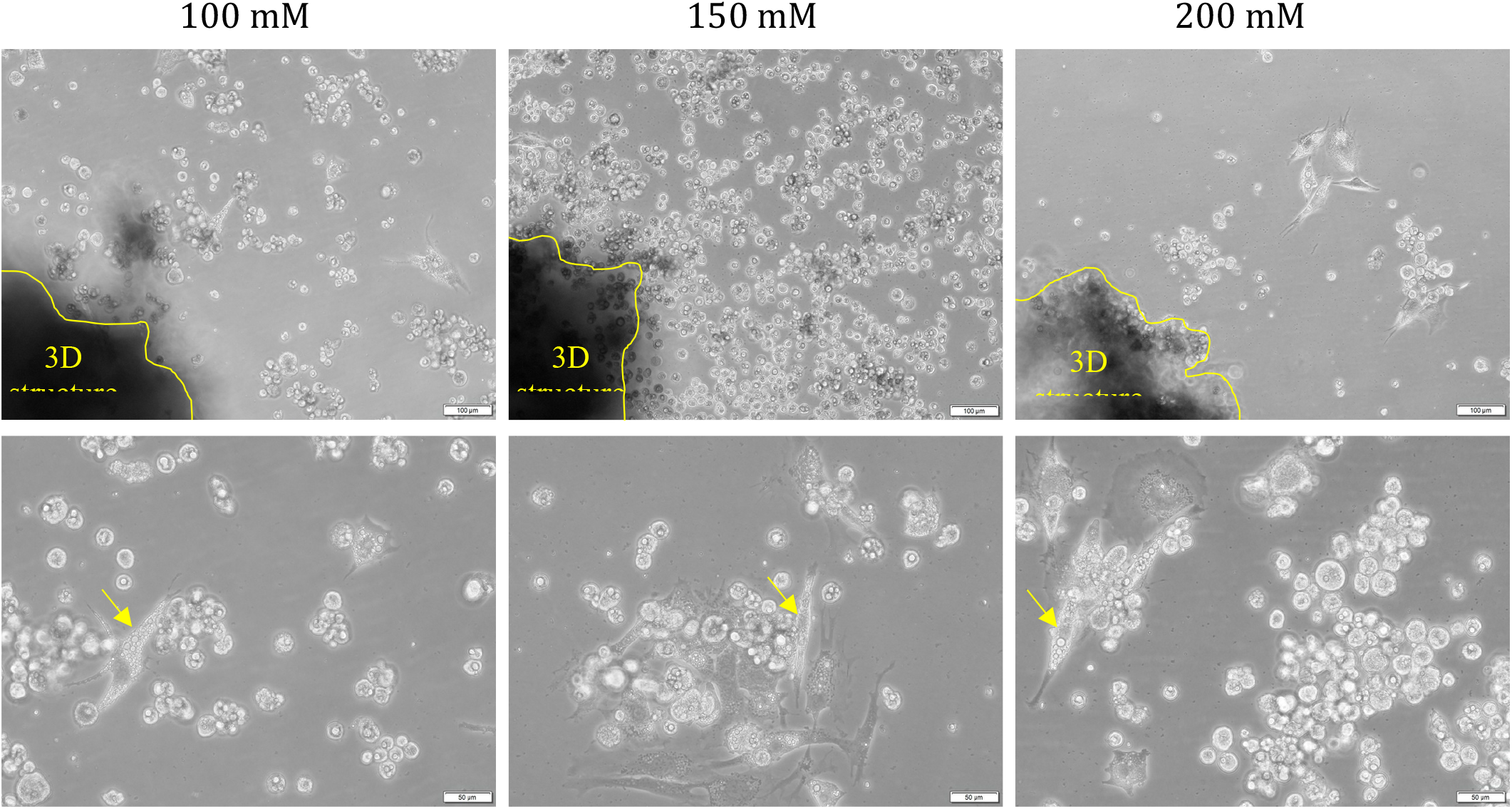
Micrographs of the adipogenesis induced 3D structures, which were formed as a result of 24-hr levitation (100, 150 and 200 mM Gd^3+^, 10^6^ cells/mL, 200 µL) and then cultured for another 24 hrs on a culture plate. The edges of the 3D structure in the image are roughly drawn with yellow lines. Arrows show examples of lipid-accumulated cells spread on the culture surface. Scale bars: 100 µm for the upper images and 50 µm for the lower images.

**Supplementary Figure 10.**
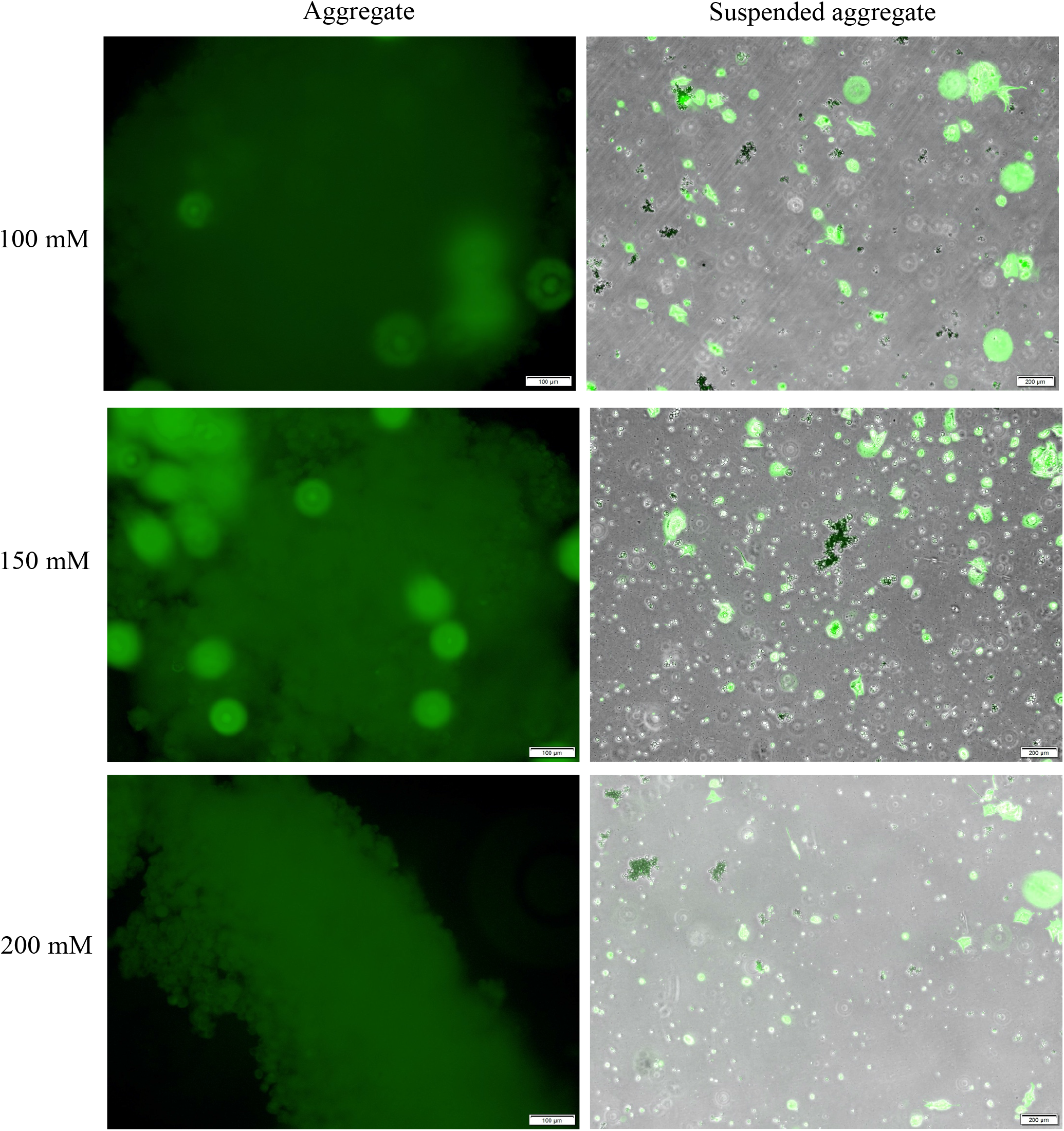
Fluorescent microscopy images of adipogenesis induced 7F2 3D structures formed with magnetic levitation and cells dissociated from the 3D structures. Cell viability was visualized by live staining (Calcein-AM). Scale bars: 100 µm for the left images and 200 µm for the right images.

**Supplementary Figure 11.**
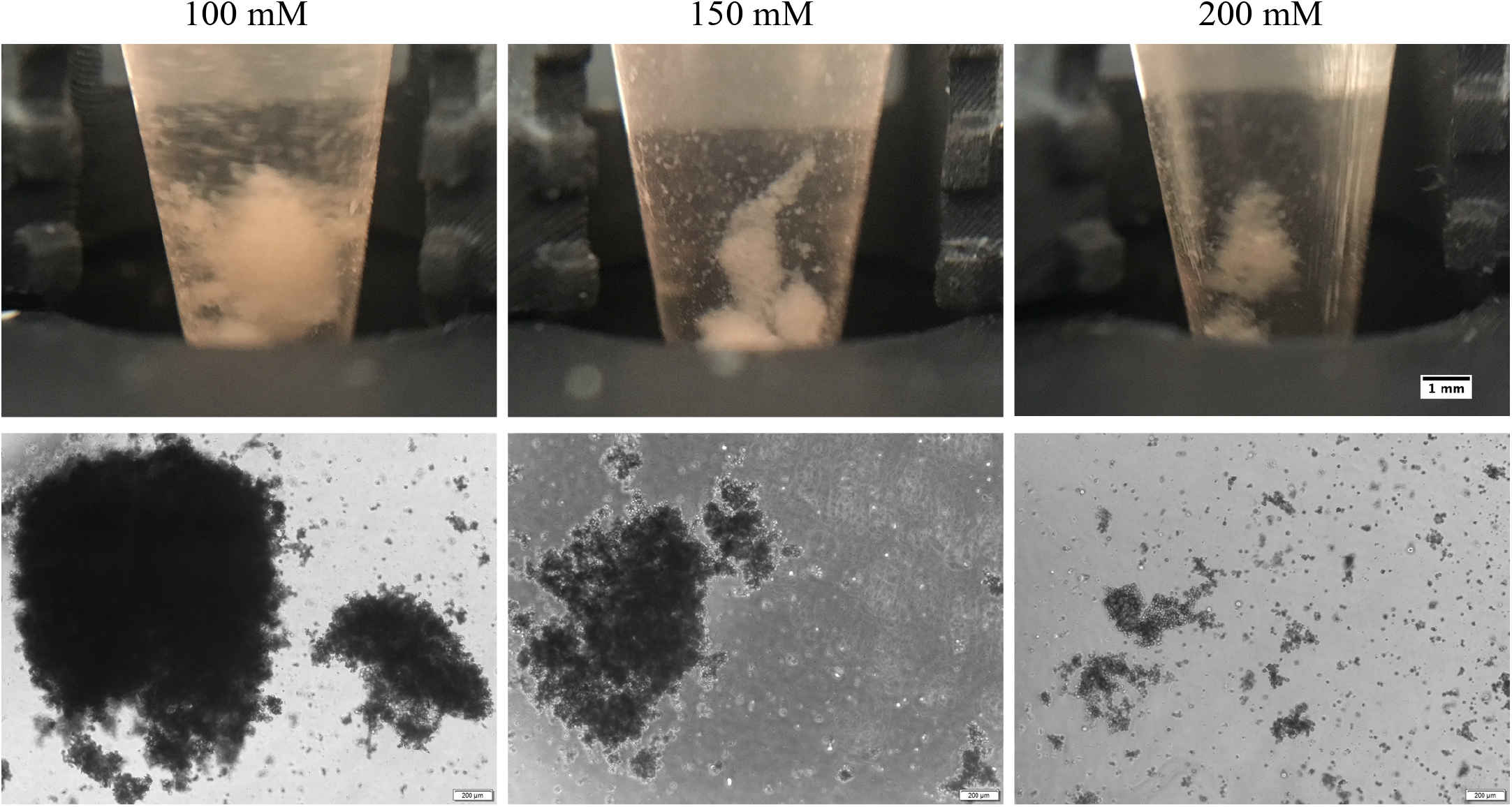
Co-levitation culture of preformed two adipogenesis induced 7F2 3D structures in medium containing 100, 150 and 200 mM Gd^3+^ in the magnetic levitation system for 24 hrs. The micrographs in the bottom row show these co-levitation products after transfer to the 2D culture dish. Each vertical unit on the 3D printed scaled piece: 1mm. Scale bars: 1 mm for culture chamber images, 200 µm for microscope images.

**Supplementary video 14**. Embedding a D1 ORL UVA 3D cluster assembled by magnetic levitation within Matrigel.

**Supplementary video 15**. Harvest of the Matrigel-embedded D1 ORL UVA 3D cluster from levitation system.

**Supplementary Figure 12.**
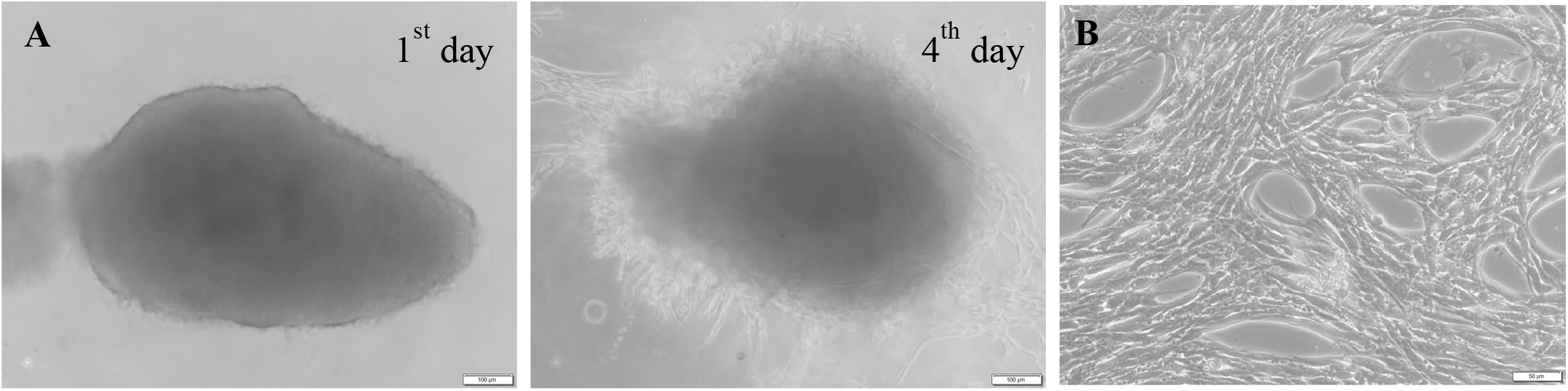
Micrographs of (A) the Matrigel-embedded 3D cluster after 1 and 4 days of culture, and (B) cells spread in Matrigel after 4 days. Scale bars: 100 µm for (A) and 50 µm for (B).

